# A Scalable Sign-Aware Multi-Omics Knowledge Graph Foundation Model for Mechanistic Drug Action and Clinical Response Predictions

**DOI:** 10.64898/2026.04.29.721775

**Authors:** Mohammadsadeq Mottaqi, Shuo Zhang, Ian Adoremos, Pengyue Zhang, Lei Xie

## Abstract

Mechanistically predicting drug action requires distinguishing activating from inhibitory interactions across broad chemical space, yet most biomedical knowledge graphs and graph neural networks (GNNs) rely on unsigned associations that obscure regulatory logic and have a limited chemical coverage. Here we present SIGMA-KG (**SIG**ned **M**ulti-omics **A**tlas **K**nowledge **G**raph) and FLASH (**F**ast **L**ightweight **A**rchitecture for **S**igned **H**eterogeneous GNN), a graph foundation model pretrained through self-supervised learning on SIGMA-KG. SIGMA-KG integrates chemogenomic perturbation, transcriptomic, proteomic, and clinical data while explicitly encoding biological polarity and directionality. FLASH preserves polarity composition across multi-hop pathways through structural balance principles, enabling scalable mechanistic reasoning. Without task-specific fine-tuning, FLASH consistently outperforms or matches nine state-of-the-art unsigned, relational, and signed graph baselines across drug mode-of-action prediction, clinical response modeling, and drug–drug interaction prediction, while substantially improving computational efficiency. FLASH further enables explainable inductive drug repurposing, achieving a 69.6% external clinical validation success rate across four complex diseases.

## 1 Introduction

Understanding the complex interactions between drugs, genes, proteins, and diseases is fundamental to drug discovery, precision medicine, and chemical safety assessments [1]. Recent advances in graph neural networks (GNNs) have enabled powerful models that capture the topological and relational structure of biomedical knowledge graphs (KGs), learning embeddings that jointly encode biological semantics and network connectivity [2]. These representations show promise for predicting drug-target interactions (DTIs) [3], drug-drug interactions (DDIs) [4], and identifying new therapeutic uses for existing drugs (drug repurposing) [5]. However, existing frameworks remain limited in their ability to capture the mechanistic logic underlying drug action. The first limitation is chemical coverage. Many widely used biomedical KGs, including Hetionet (47,031 nodes; 2.2 million edges) [6], PrimeKG (129,375 nodes; 4 million edges) [7], and BioPathNet(32,000 edges) [8], focus primarily on only a few thousand approved or well-annotated compounds, thereby under-representing the much larger chemical space relevant to early-stage discovery. The second limitation is that all of these KGs encode interactions as unsigned relations without explicit effect polarity, thereby conflating activation with inhibition. Signed biological databases address polarity in more restricted settings: SIGNOR 2.0 encodes ∼33,000 activation or inhibition edges across proteins, RNA, and metabolites [9], whereas OmniPath [10] and TRRUST v2 [11] cover additional single-modality regulatory layers. Although these resources preserve directional signs, +1 for activation/up-regulation and −1 for inhibition/down-regulation, they are not integrated into a unified, multi-omics atlas that simultaneously connects chemical perturbations, molecular responses, and clinical phenotypes in a form suitable for translational modeling (see Supplemental Information).

This missing polarity information is consequential because many biological signaling and regulatory processes are inherently sign-dependent. Within these signaling networks, the functional polarity of a multi-hop pathway is governed by the composition of constituent interactions, where the net regulatory effect is determined by the multiplicative product of signs along the path [12, 13]. Consistent with this view, empirical studies of gene-gene interaction networks have shown that balanced triads are significantly overrepresented, whereas unbalanced triads are underrepresented relative to randomized controls; this suggests that interaction signs in biological networks are organized in a manner consistent with structural balance principles [14, 15]. Together, these observations support the use of sign-consistent path composition as a biologically grounded inductive bias for modeling net mechanistic effects in biological systems. The practical importance of considering polarity is illustrated by vemurafenib, a BRAF inhibitor approved for melanoma. In unsigned KGs, its proximity to the ERK signaling cascade could suggest therapeutic benefit; yet in *KRAS* -mutant cells, the same drug can paradoxically hyperactivate ERK and promote tumor growth [16]. Such context-dependent sign reversals are difficult to capture with unsigned proximity-based models.

While relational (heterogeneous) GNNs (rGNNs) assign distinct embeddings to different edge types, they still represent an inhibition or activation interaction as a categorical label rather than an algebraic operator, and therefore cannot model signal transduction along multi-hop paths (e.g., *A* −→ *B* −−→ *C* =⇒ *A* −+→ *C*) [13].

We define this conceptual failure as *polarity drift* : the systematic divergence between a model’s predicted functional direction and the ground-truth sign-product of a regulatory pathway. This divergence compounds as path length increases, necessitating signed Graph Neural Networks (sGNNs) that explicitly integrate edge polarity into the message-passing framework. By grounding representation learning in structural balance principles, sGNNs facilitate robust multi-hop reasoning that preserves the integrity of long-range signaling logic [17].

Large language models (LLMs) provide a complementary source of biomedical knowledge through text, but they do not explicitly model sign-consistent propagation over graph topology. They retrieve pairwise associations from the literature, yet they are not designed to reliably propagate the product of signed directed edges across multi-hop regulatory paths, especially when such paths are not explicitly described in training corpora [18]. For this reason, graph-based learning with explicit sign semantics remains a distinct and necessary modeling framework for mechanistic prediction in biomedicine [19].

Recent studies applying sGNNs to biological graphs support the promise of sign-aware modeling, while also highlighting current limitations. SIGDR [20] uses sign-aware contrastive learning to model drug-disease associations, and CSGDN [21] applies signed graph diffusion to predict crop gene-phenotype associations. These studies demonstrate the utility of signed graph learning in biology, but they adopt supervised learning objectives, precluding self-supervised pre-training on unlabeled network topology, and operate on single-modality, small-scale graphs (*<* 20, 000 edges) that are not scalable to the multi-omics, million-edge setting required for comprehensive drug action modeling.

In parallel, graph foundation models have emerged as a promising direction for biomedicine by showing that pre-training on large KGs can yield transferable representations for drug repurposing, DDI prediction, and gene function annotation [8, 22]. TxGNN [5] is the most directly relevant predecessor: trained on a relational KG comprising 17,080 diseases and 7,957 drug candidates [7], it performs indication and contraindication ranking, substantially improving over earlier approaches. However, TxGNN relies on supervised metric-learning objectives defined over labeled drug-disease associations, which may limit generalization to diseases with scarce or no known therapeutic examples. Additionally, its underlying KG does not encode edge polarity and covers no early-stage small molecules, preventing sign-consistent multi-hop reasoning. BioPathNet [8] advances link prediction via path-based neural reasoning on an unsigned KG, but it is trained in a fully supervised manner. PT-KGNN [22] introduces self-supervised pre-training on unsigned biomedical KGs and demonstrates scaling with KG size, yet it applies homogeneous aggregation that discards relational and sign semantics. Collectively, these models expose a persistent gap: no existing framework combines *self-supervised* pre-training with explicit directed edge-sign semantics in a large-scale, multi-omics atlas.

We therefore hypothesize that explicitly encoding directed edge polarity in a large-scale multi-omics KG, and using this graph to pre-train a sign-aware GNN in a self-supervised manner, would produce more accurate and mechanistically interpretable predictions of drug actions than unsigned homogeneous or relational alternatives. To test this hypothesis, we constructed SIGMA-KG, a signed and directed multi-omics atlas of ∼127,000 nodes, including 76,009 small molecules and 3.8 million signed edges integrated from 16 heterogeneous biomedical resources. We further developed FLASH, a computationally efficient signed heterogeneous GNN that enables the pre-training of large-scale signed KG in a self-supervised manner to learn transferable, balance-aware node representations without requiring task-specific supervision.

To evaluate the advantages of sign-aware representation learning, we benchmarked FLASH on three critical tasks spanning molecular to clinical scales: : target-specific mode-of-action (Ts-MoA), drug-induced clinical response (DCR), and DDI predictions. Across these benchmarks, FLASH matched or exceeded nine SOTA unsigned, relational, and signed baselines while maintaining high computational efficiency. Furthermore, we illustrated the model’s translational utility through explainable inductive drug repurposing, where high-confidence therapeutic candidates for complex diseases are recovered via biologically coherent, multi-hop signed paths. These results establish SIGMA-KG and FLASH as a scalable foundation for sign-aware, mechanistically grounded predictive modeling in drug discovery and clinical pharmacology.

## 2 Results

### 2.1 SIGMA-KG and FLASH establish a scalable sign-aware foundation-model framework for drug discovery

Here we introduce a scalable foundation-model framework for mechanistic drug discovery and drug repurposing, centered on SIGMA-KG and FLASH. Unlike conventional unsigned GNNs that overlook the polarity of biological interactions, our framework explicitly models signed biological effects and leverages structural balance principles to support sign-aware learning across molecular, clinical, and systems-level pharmacological settings.

We constructed SIGMA-KG, a large-scale, signed, multi-relational atlas that bridges molecular perturbations and clinical outcomes **(Fig. 1a; Supplementary Table 1)**. Specifically, we harmonized 16 large-scale resources spanning chemical genomics, transcriptomics, proteomics, and clinical phenotypes into a unified, signed, multi-layer KG. The resulting atlas comprises 127,422 nodes across four primary biomedical entity types (compounds, genes, proteins, and diseases) and 3.88 million signed edges **(Table 1; Supplementary Table 2)**. These edges encode explicit biological polarities (a signed label per edge, +1 or −1), including activation versus inhibition, up-regulation versus down-regulation, and therapeutic indication versus adverse contraindication, thereby providing a mechanistically rich substrate for sign-aware modeling **(Supplementary Figs. 1a and 1b)**.

**Fig. 1.**
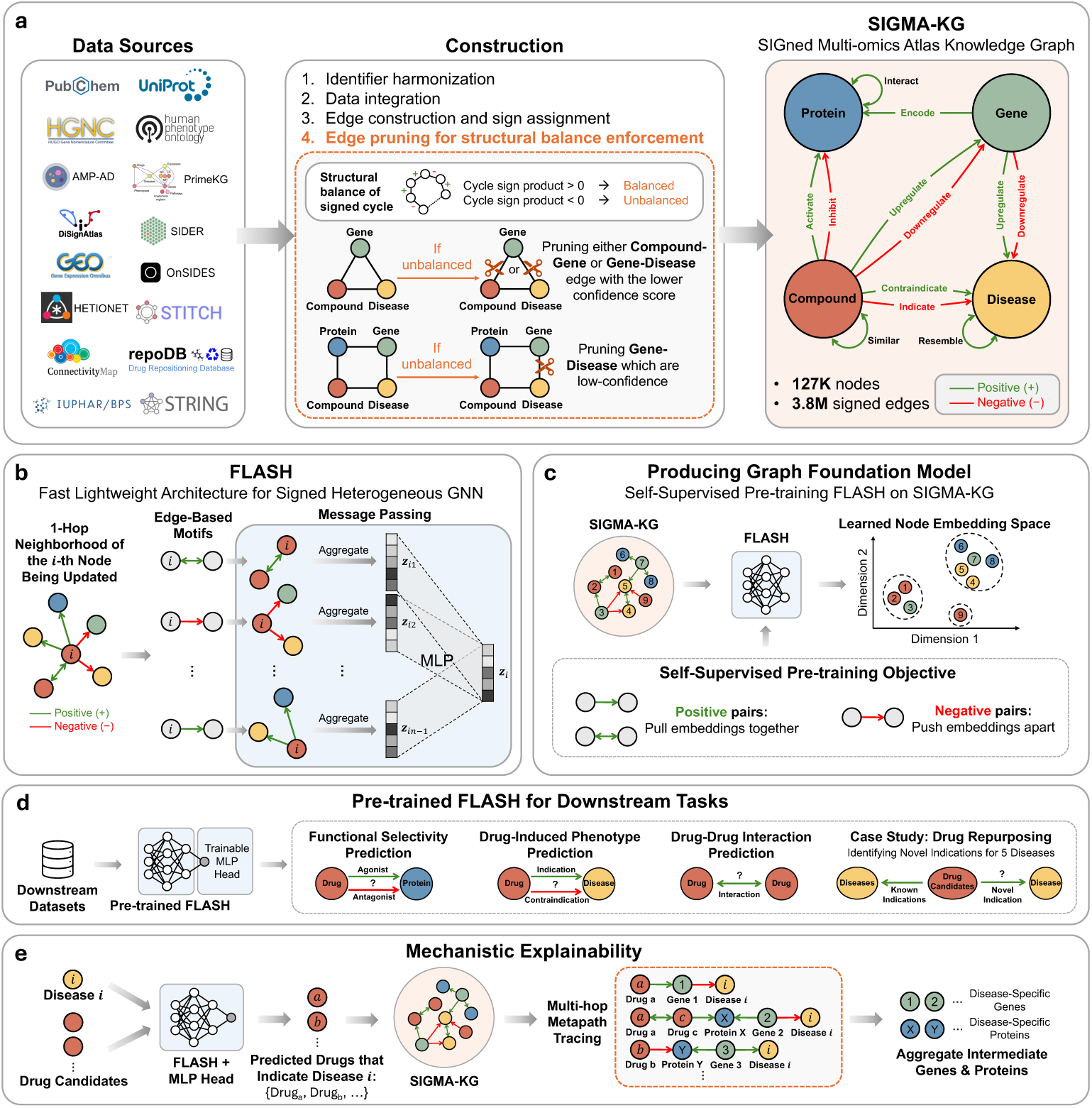
Sign-aware multi-omics graph foundation model for mechanistic drug action prediction across molecular-to-clinical scales. a, Construction of SIGMA-KG from harmonized identifiers and integrated multi-omics and clinical sources, forming a heterogeneous directed graph of compounds, genes, proteins, and diseases. Edge signs encode mechanistic polarity (positive or negative), including activation/inhibition, upregulation/downregulation, and indication/contraindication. Structural balance is enforced by pruning low-confidence edges involved in unbalanced signed cycles with sign product *<* 0. Pruning preferentially removes lower-confidence compound-gene or gene-disease edges. The resulting graph contains ∼127K nodes and ∼3.8M signed edges. **b**, FLASH encodes each node using 1-hop neighborhood aggregation conditioned on edge-based signed motifs. Separate message-passing channels for positive and negative signed edges preserve transitive sign composition along multi-hop paths and mitigate sign mixing during aggregation. A multi-layer perceptron (MLP) produces the final embedding *z* for each node. **c**, Self-supervised pre-training of FLASH on the global topology of SIGMA-KG using sign-aware objectives to learn transferable graph representations. **d**, The frozen embeddings from the pre-trained foundation model is evaluated across downstream pharmacological tasks, including target-specific mode-of-action prediction (agonist/antagonist), drug-induced clinical response prediction (indication/contraindication), drug-drug interaction prediction, and a drug repurposing case study across four diseases. **e**, Mechanistic explainability via multi-hop signed metapath tracing in SIGMA-KG, aggregating intermediate disease-linked genes and proteins to support candidate drug-disease predictions.

**Table 1.** Statistical details of SIGMA-KG. SIGMA-KG comprises 127,422 nodes and 3,887,829 signed edges connecting four biomedical entity types. Percentages (%) are relative to the total number of nodes or edges, respectively. Where multiple relation types exist between the same source and target entity types, they are listed together in a single row with their corresponding sign labels.

| Entity type | Description | No. of nodes | % of all nodes |
| --- | --- | --- | --- |
| Gene | Human protein-coding genes (HGNC gene symbols) | 19,245 | 15.1% |
| Protein | Human proteins (UniProt identifiers) | 18,609 | 14.6% |
| Compound | Small molecules and drug-like chemicals (PubChem CID) | 76,009 | 59.7% |
| Disease | Human diseases and clinical phenotypes (Human Phenotype Ontology (HPO)) | 13,559 | 10.6% |
| **Total** |  | **127,422** | **100%** |

(b) Edge statistics by biomedical entity pair.
| Source entity | Target entity | Relation type(s) with sign | No. of edges | % of all edges |
| --- | --- | --- | --- | --- |
| Gene | Disease | Up-regulated in (+1); Down-regulated in (−1) | 1,489,308 | 38.3% |
| Chemical | Gene | Up-regulates (+1); Down-regulates (−1) | 1,322,830 | 34.0% |
| Compound | Compound | Similar to (+1) | 581,872 | 15.0% |
| Protein | Protein | Interacts with (+1) | 251,762 | 6.5% |
| Compound | Disease | Contraindication (+1); Indication (−1) | 159,957 | 4.1% |
| Disease | Disease | Resembles (+1) | 49,052 | 1.3% |
| Gene | Protein | Encodes (+1) | 19,337 | 0.5% |
| Compound | Protein | Activates (+1); Inhibits (−1) | 13,711 | 0.4% |
| **Total** |  |  | **3,887,829** | **100%** |

SIGMA-KG exhibits high cross-modal density, with an average node degree of 30.5, enabling the tracing of continuous mechanistic chains from molecular perturbations to clinical phenotypes. In total, the atlas links 22,862 compounds directly to down-stream transcriptomic signatures and proteomic targets, which are further connected to a unified disease landscape of 13,559 clinical phenotypes. This organization substantially expands the mechanistically reachable therapeutic space. Although only 3.8% of compounds (2,869/76,009) are directly linked to diseases through recorded clinical associations, the proportion of compounds connected to at least one disease phenotype increases to 28.5% (21,649/76,009) within two hops and 47.2% (35,905/76,009) within three hops. These results show that SIGMA-KG is not simply a collection of disjoint datasets, but a dense, searchable multi-omics map through which biological effects can be propagated across scales. We further examined the signed topology of SIGMA-KG through the lens of Heider’s structural balance theory [14] **(Supplementary Fig. 1c)**. Across closed triads in the atlas, 99.6% were balanced, indicating that the signed graph structure is highly non-random and largely consistent with structural balance principles **(Supplementary Fig. 1d)**. This high degree of triadic consistency supports the use of sign-aware graph learning as a biologically grounded inductive bias for modeling multi-hop effects in the network.

Leveraging SIGMA-KG, we developed FLASH, a scalable signed heterogeneous GNN designed for atlas-scale representation learning **(Fig. 1b)**. Unlike standard relational GNNs, FLASH utilizes a motif-aware attention mechanism to encode seven biologically informed signed patterns, explicitly preserving signaling polarity and the ’double negative’ logic where inhibiting an inhibitor results in net activation. In self-supervised pre-training, FLASH uses observed edges and their signs in SIGMA-KG as supervision to optimize the link sign product loss, a sign-aware structural objective that encourages similar embeddings for positively connected nodes and dissimilar embeddings for negatively connected nodes (Methods). This process yields robust, transferable node embeddings that capture both topological context and sign-aware biomedical semantics **(Fig. 1c)**.

FLASH is computationally efficient while retaining the expressive capacity required for multi-hop sign-aware reasoning. Compared with existing signed GNNs, the framework achieves a substantial reduction in training time, while producing a general-purpose graph foundation model that can be applied across diverse downstream pharmacological tasks. In the following sections, we evaluate this foundation model across three biological scales: Ts-MoA prediction at molecular level, DCR prediction at clinical level, and DDI prediction at systems level, followed by explainable inductive drug repurposing analyses **(Fig. 1d and 1e)**.

To evaluate the predictive utility of the pretrained node embeddings, we implemented a standardized benchmarking protocol comparing FLASH against nine unsupervised GNN baselines. For molecular-level (Ts-MoA) and clinical-level (DCR) tasks, we employed an internal validation strategy using a randomized edge-split protocol. To prevent data leakage, SIGMA-KG was partitioned into three independent sets of held-out evaluation edges; the models were then pretrained in a self-supervised manner on the remaining graph topology to generate 256-dimensional node embeddings. These frozen embeddings were subsequently evaluated via a Multi-Layer Perceptron (MLP) head—trained on the internal training edges—to perform binary and three-class edge-sign prediction on the held-out sets.

In contrast, systems-level DDI prediction followed an external validation strategy. Models were pretrained on the complete SIGMA-KG atlas, and the resulting compound embeddings were directly assessed on two independent DDI benchmark datasets: DrugBank [23] and MUDI [24]. To ensure a controlled comparison, all models were initialized with identical 256-dimensional node features derived from truncated singular value decomposition (SVD). Evaluation across all tasks utilized standardized MLP architectures and hyperparameter tuning protocols (Methods). By standardizing architectural components and feature initialization, this experimental design isolates the specific predictive contribution of the signed multi-relational inductive bias from confounding factors. This enables a rigorous head-to-head comparison between FLASH and existing baselines while providing a clear basis for demonstrating the systematic performance advantage of sign-aware modeling over standard relational and unsigned graph architectures.

### 2.2 FLASH enables robust target-specific mode-of-action prediction

We first evaluated the capacity of the SIGMA-KG-pretrained foundation model to accurately predict Ts-MoA, a critical prerequisite for understanding the downstream functional consequences of drug-target engagement [3]. This task is central to pharmacology as it distinguishes whether a protein ligand acts as an agonist (target activator) or an antagonist (target inhibitor), a distinction that conventional GNNs often overlook by treating DTIs as unsigned proximities. We formulated Ts-MoA prediction as an edge-sign prediction task in both binary (activation, +1 vs. inhibition, −1) and three-class classification settings, with the latter incorporated a 5:1 ratio of non-existent associations (0) to labeled edges [25] (Methods). These evaluations were conducted on held-out datasets of Compound–Protein interactions that were entirely unseen during model pre-training, ensuring a rigorous assessment of the model’s ability to generalize to novel molecular mechanisms.

Our analysis showed that signed GNN architectures overall outperformed unsigned and relational baselines in Ts-MoA prediction **(Supplementary Tables 3and 4)**, with FLASH achieving SOTA or comparable performance across both binary and three-class settings **(Fig. 2 and Supplementary Fig. 2)**. In the binary classification setting **(Fig. 2a-c, Supplementary Table 5)**, the top signed baselines (FLASH and SNEA) significantly outperformed the top unsigned baseline (GraphSAGE) in both Accuracy (3.63% higher, *p* = 0.014, unpaired Welch’s t-test) and Macro-F1 (3.51% higher, *p* = 0.020). Within this setting, FLASH achieved the highest overall Accuracy (0.8990 ± 0.0143) and showed performance statistically indistinguishable from the top-performing signed baseline (SGCN) in AUROC (*p* = 0.980) and Macro-F1 (*p* = 0.836; Supplementary Table 4.

**Fig. 2.**
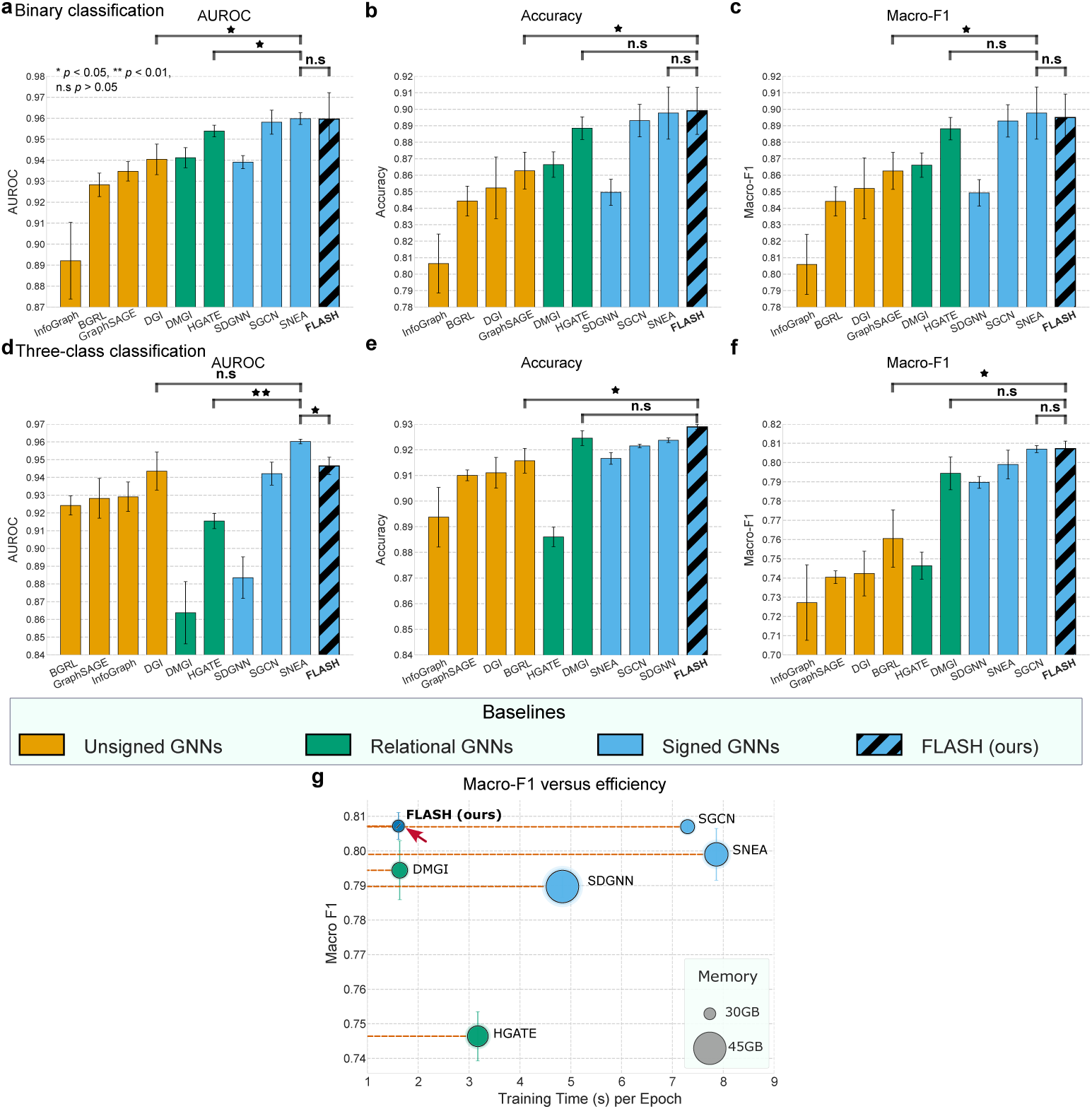
Performance comparison of FLASH with unsigned, relational, and signed graph neural network (GNN) baselines for predicting target-specific mode-of-action. **a–c**, Binary classification performance (activation or inhibition). Comparison across baselines using (a) AUROC, (b) accuracy, and (c) macro-F1 score. **d–f**, Three-class classification performance (activation, inhibition, or no interaction) evaluated using (d) AUROC, (e) accuracy, and (f) macro-F1 score. Bars represent the mean across three independent runs with random train–test splits, with error bars indicating the standard deviation. Statistical significance was assessed using two-tailed paired t-tests. FLASH was compared against the overall top-performing baseline or, in instances where FLASH established the state-of-the-art, the next-best performing model. Additionally, the leading signed baseline was compared against the top relational and unsigned alternatives to quantify the predictive gains afforded by explicit edge polarity in SIGMA-KG dataset. Significance annotations are shown above brackets (∗*p <* 0.05, ∗ ∗ *p <* 0.01, n.s., not significant). Complete P-values are reported in Supplementary Tables 17 and 18, and full metric results for binary and three-class tasks are provided in Supplementary Tables 5 and 6. **g**, Accuracy–efficiency trade-off. Macro-F1 score for three-class classification plotted against training time per epoch for the top-performing signed and heterogeneous GNN baselines, illustrating the accuracy–efficiency trade-off. Marker size indicates approximate memory usage during training. FLASH achieves competitive or superior predictive performance while maintaining substantially lower training time compared with other top-performing signed GNN architectures.

Similarly, in the more challenging three-class setting **(Fig. 2d-f), Supplementary Table 6**, which requires joint discrimination of activation, inhibition, and the absence of an interaction, signed baselines demonstrated a substantial advantage in interaction detection and polarity classification, with FLASH significantly outperform-ing the best unsigned baseline (BGRL) in both Accuracy (1.32% higher, *p* = 0.0183) and Macro-F1 (4.67% higher, *p* = 0.0130). Furthermore, in the three-class setting, the top signed baseline (SNEA) significantly outperformed the top relational model (HGATE) by 4.48% in Macro AUROC (*p* ≤ 0.001), highlighting the advantage of signed architectures in distinguishing true functional interactions from biological noise. Although SNEA maintained a statistically significant lead in three-class Macro AUROC (*p* = 0.032), FLASH’s superior F1 scores for both inhibition and non-existent edges suggest a more balanced ability to capture functional polarity while limiting false positives **(Supplementary Fig. 2**.

Beyond its predictive accuracy, FLASH demonstrates superior computational scalability, a prerequisite for atlas-scale inference. Quantitatively, FLASH achieved SOTA Macro-F1 scores while requiring 4.5-fold less training time per epoch and substantially lower memory overhead compared to the next-best signed baseline, SGCN **(Fig. 2g)**. These results establish FLASH as a signed graph foundation model capable of high-fidelity Ts-MoA inference at atlas scale while maintaining computational efficiency.

### 2.3 FLASH enables robust drug-induced clinical response prediction

Accurate DCR prediction o is essential for prioritizing therapeutic indications while mitigating adverse contraindications [5]. Distinguishing between these outcomes is a primary challenge in drug development, exacerbated by a severe class imbalance where documented contraindications (in SIGMA-KG) significantly outnumber therapeutic indications (∼12.4:1). We evaluated the capacity of FLASH using its frozen embeddings to navigate this imbalanced landscape by predicting DCR, formulated as an edge-sign prediction task in both binary (indication, −1 vs. contraindication, +1) and three-class classification settings, with the latter incorporating a 5:1 ratio of non-existent associations (0) to labeled edges to serve as a rigorous proxy for real-world inductive drug repurposing [25] (Methods). These evaluations were performed on held-out datasets of Compound–Disease interactions entirely excluded from pre-training, testing the model’s ability to recover sparse, high-stakes therapeutic signals from the vast landscape of clinical associations.

Our comparative analysis revealed that signed baselines consistently outperformed top-performing unsigned and relational GNNs **(Supplementary Tables 3 and 4)**. In the binary setting **(Fig. 3a-c)**, the top-performing signed models, FLASH and SDGNN, exhibited a significant performance advantage in both Accuracy and Macro-F1, with FLASH outperforming the top unsigned baseline (BGRL; *p* = 0.002) and the leading relational baseline (HGATE; *p* ≤ 0.001) in Accuracy. This performance gap widened in the more demanding three-class setting **(Fig. 3d-f)**, where the top signed model (SGCN) outperformed the leading unsigned model (DGI) by 11.46% in Macro-F1 (*p* = 0.003) and the top-performing relational model (HGATE) by 3.32% (*p* = 0.017) While unsigned models such as DGI achieved high AUPRC (0.9995) in the binary setting, their inability to model directionality led to a substantial failure in identifying minority-class therapeutic indications in the three-class setting (F1_Indication_ ≤ 0.51), whereas signed models such as FLASH more effectively prioritized these relationships (F1_Indication_ = 0.81; **Supplementary Figs. 2b and 2d**).

**Fig. 3.**
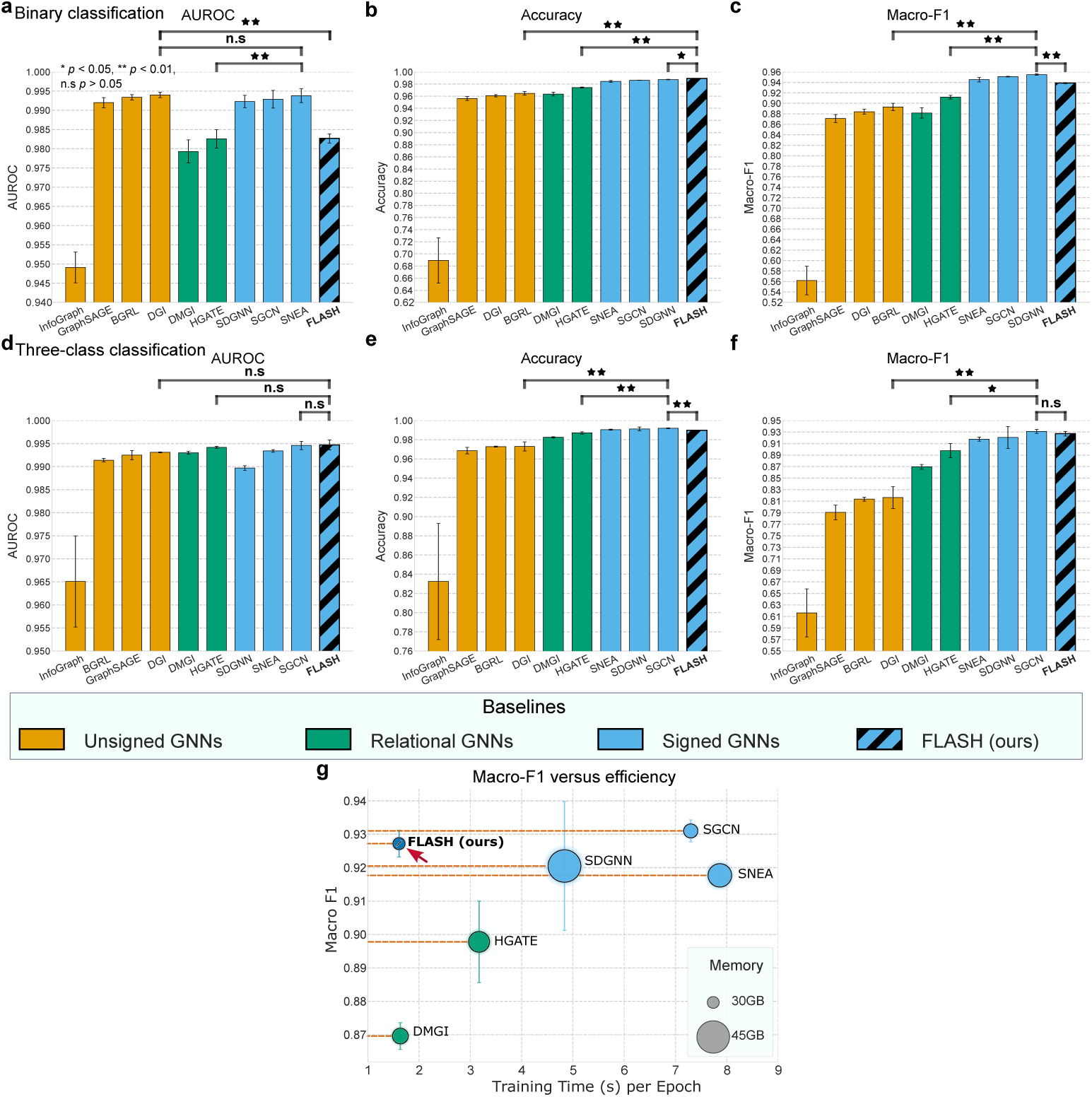
Performance comparison of FLASH with unsigned, relational, and signed graph neural network (GNN) baselines for predicting DCR relationships. **a–c**, Binary classification performance. Comparison across baselines using (a) AUROC, (b) accuracy, and (c) macro-F1 score. **d–f**, Three-class classification performance (indication, contraindication, and no association) evaluated using (d) AUROC, (e) accuracy, and (f) macro-F1 score. Bars represent the mean across three independent runs with random train–test splits, with error bars indicating the standard deviation. Statistical significance was assessed using two-tailed paired t-tests. FLASH was compared against the overall top-performing baseline or, in instances where FLASH established the state-of-the-art, the next-best performing model. Additionally, the leading signed baseline was compared against the top relational and unsigned alternatives to quantify the predictive gains afforded by explicit edge polarity in SIGMA-KG dataset. Significance annotations are shown above brackets (∗*p <* 0.05, ∗ ∗ *p <* 0.01, n.s., not significant). Complete P-values are reported in Supplementary Table A1, and full metric results for binary and three-class tasks are provided in Supplementary Tables 7 and 8. **g**, Accuracy–efficiency trade-off. Macro-F1 score for three-class classification plotted against training time per epoch for the top-performing signed and heterogeneous GNN baselines. Marker size reflects approximate GPU memory usage during training. FLASH achieves competitive or superior predictive performance while maintaining substantially lower training time compared with other high-performing signed GNN architectures.

Within this overall pattern, FLASH showed strong performance in the binary setting **(Supplementary Table 7)**. In detail, FLASH attained the highest over-all Accuracy (0.9893 ± 0.0003), significantly outperforming the runner-up baseline (SDGNN; *p* = 0.008) while establishing a 1.51% lead over the leading relational baseline (HGATE; *p* ≤ 0.001).

Beyond binary classification, FLASH also maintained strong performance in the more challenging three-class setting which includes non-existent associations **(Supplementary Table 7)**. FLASH demonstrated strong discriminative power, achieving the highest Macro AUROC (0.9947 ± 0.0011). While SGCN showed a marginal statistical advantage in Accuracy (Δ = 0.22%, *p* = 0.002), FLASH maintained comparable Macro-F1 performance (*p* = 0.275), and achieved a 4.5-fold reduction in training time per epoch **(Fig. 3g)**. This capacity to maintain high Macro AUROC and F1 scores under severe class imbalance, while remaining computationally efficient, provides a strong foundation for our subsequent therapeutic indication prediction task, ensuring that the predicted drug repurposing candidates are prioritized based on robust, sign-aware structural evidence.

### 2.4 FLASH significantly improves drug-drug interaction prediction across both warm- and cold-start scenarios

Accurate DDI prediction is a cornerstone of clinical safety and rational polypharmacy [26]. Yet this application remains a formidable challenge, particularly when models must generalize to novel therapeutic contexts where interaction data are sparse. To evaluate the pharmacological utility of the representations learned by our foundation model, we assessed its capacity to predict DDIs. Crucially, within SIGMA-KG, compound-compound edges are derived exclusively from molecular structural similarity, meaning that the foundation model receives no explicit pharmacological or clinical DDI knowledge during pre-training. The DDI benchmark thus evaluates whether FLASH can synthesize molecular structure with multi-omics signaling to predict complex interactions. This capability is critical for capturing DDIs that are not encoded within chemical structures alone, but instead emerge through the network-mediated propagation of biological changes across the atlas.

We benchmarked our framework using two external DDI benchmark datasets, DrugBank (version 5.1) [23] and MUDI [24]. DrugBank dataset contains clinically documented interactions curated from regulatory labels and literature, while MUDI dataset incorporates pharmacologically derived synergistic and antagonistic effects. Performance was evaluated via a binary classification task—distinguishing the presence or absence of interactions—under two distinct evaluation scenarios: transductive (warm-start) prediction based on random pair splitting, and inductive (cold-start) prediction using a drug-disjoint split to evaluate generalization to novel compounds **(Fig. 4 and Supplementary Fig. 3)**. While the transductive scenario measures the model’s ability to interpolate within known drug spaces, the inductive scenario provides a more stringent, real-world assessment by requiring the model to predict interactions for compounds entirely absent from the training set (Methods).

**Fig. 4.**
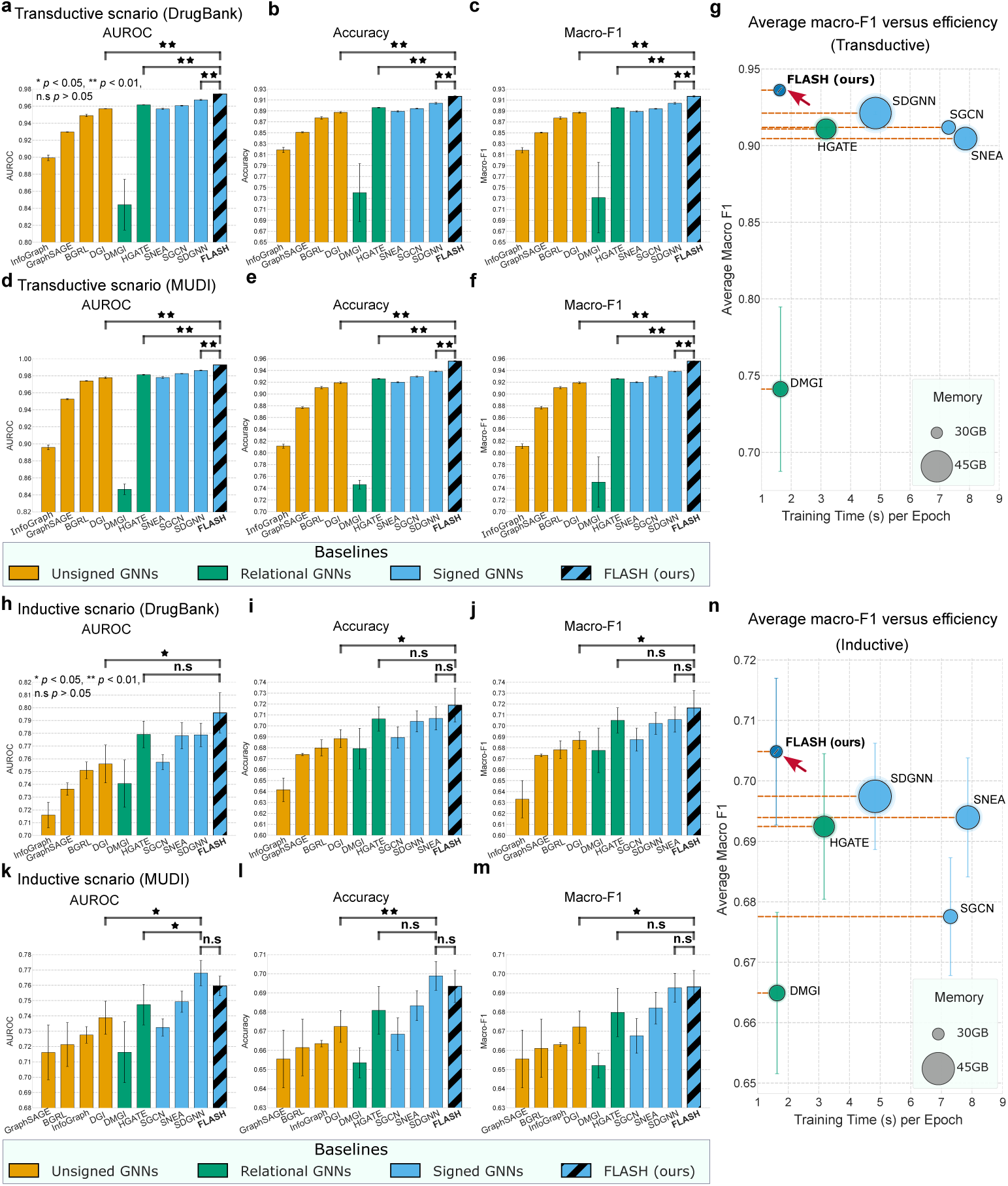
Performance comparison of FLASH with unsigned, relational, and signed graph neural network (GNN) baselines for predicting drug–drug interactions. Model performance was evaluated using AUROC, accuracy, and macro-F1. Efficiency was assessed by plotting average macro-F1 against training time per epoch, with marker size indicating GPU memory usage during training. **a-c**, Transductive prediction on DrugBank. **d-f**, Transductive prediction on MUDI. **g**, Efficiency analysis (transductive). **h-j**, Inductive prediction on DrugBank. **k-m**, Inductive prediction on MUDI. **n**, Efficiency analysis (inductive). Bars represent the mean across three independent runs with random splits, with error bars indicating the standard deviation. Statistical significance was assessed using two-tailed paired t-tests. FLASH was compared against the overall top-performing baseline or, in instances where FLASH established the state-of-the-art, the next-best performing model. The leading signed baseline was compared against the top relational and unsigned alternatives to quantify the predictive gains afforded by explicit edge polarity in SIGMA-KG. Significance annotations above brackets indicate (∗*p <* 0.05, ∗ ∗ *p <* 0.01, n.s., not significant). Complete P-values are reported in Supplementary Tables 17 and 18. ^13^

#### 2.4.1 Transductive DDI prediction scenario

In the transductive setting, the results showed that signed models incorporating edge polarity consistently outperform both unsigned and relational baselines **(Supplementary Tables 3 and 4)**. Across both DrugBank and MUDI benchmarks, the node representations learned by FLASH significantly outperformed the leading unsigned (DGI) and relational (HGATE) models in all evaluated metrics **(Fig. 4a-f, Supplementary Tables 9 and 10)**. Specifically, on DrugBank, FLASH significantly improved AUROC and Macro-F1 by 1.72% and 2.94% (both *p <* 0.0001) respectively against the top unsigned model (DGI), and by 1.26% (*p <* 0.0001) and 2.07% (*p* = 0.0002) against leading relational model, HGATE. These margins were even more pronounced on the MUDI dataset, where FLASH exceeded the Macro-F1 of DGI and HGATE by 3.66% and 2.99%, respectively (both *p* = 0.0001).

In this systems-level evaluation, FLASH established new state-of-the-art (SOTA) performance across multiple benchmarks. On the DrugBank dataset **(Fig. 4a-c)**, FLASH significantly outperformed the leading signed baseline (SDGNN) in AUROC (0.9742 ± 0.0004), AUPRC (0.9735 ± 0.0005), and Macro-F1 (0.9166 ± 0.0015; all three *p <* 0.001). This superior predictive capability was sustained on the MUDI dataset (**Fig. 4d-f**), where FLASH achieved a Macro-F1 of 0.9557 ± 0.0005, a significant 1.75% improvement over SDGNN (*p <* 0.001). Notably, these performance gains were achieved with high computational efficiency; FLASH required 3-fold less training time per epoch and significantly lower memory overhead compared to SDGNN, underscoring its practical utility for atlas-scale biomedical modeling **(Fig. 4g)**.

#### 2.4.2 Inductive DDI prediction scenario

We next assessed whether the inductive biases of signed graph pre-training enhance the model’s generalization to novel compounds. To ensure a rigorous evaluation, we implemented a drug-disjoint partitioning strategy (splitting the unique set of compounds into training, validation, and test sets) such that every test-set interaction involved at least one compound entirely unseen during the classifier’s training phase. Although predictive performance across all architectures decreased in this more challenging cold-start scenario, signed and relational models exhibited superior robustness compared to unsigned baselines, maintaining significantly higher generalizability **(Fig. 4h-m, Supplementary Tables 3and 4)**.

On the DrugBank dataset **(Fig. 4h-j; Supplementary Table 11)**, FLASH out-performed all baselines, including the leading relational model (HGATE), achieving an AUROC of 0.7960 ± 0.0157 (*p* = 0.20) and an AUPRC of 0.7971 ± 0.0192 (*p* = 0.18).

While the performance gap between the top models narrowed in this high-difficulty setting, FLASH remained among the leading methods across metrics. On MUDI dataset **(Fig. 4k-m; Supplementary Table 12)**, FLASH achieved the highest Macro-F1 (0.6932 ± 0.0085; **(Fig. 4n)**) while showing comparable performance to the top baseline (SDGNN) in AUPRC (*p* = 0.344). Although SDGNN showed a marginal lead in Accuracy (0.6989 vs 0.6934, *p* = 0.445), the performance of FLASH remains highly competitive.

Taken together, these results show that FLASH offers a favorable balance between predictive performance and computational efficiency for large-scale biomedical applications. Across all scenarios, FLASH achieved comparable or superior Macro-F1 scores while benefiting from the reduction in training time and memory overhead compared to other signed and relational models **(Fig. 4g and Fig. 4n)**. Systematic bench-marking across varying graph scales further confirms that FLASH maintains linear scalability and a significantly reduced memory footprint, as detailed in Supplementary Information and **Supplementary Fig. 4**. Collectively, these findings suggest that by leveraging the signed multi-omics structure of SIGMA-KG, FLASH learns latent representations that reflect functional and mechanistic similarity rather than molecular structure alone, enabling robust sign-aware prediction of drug actions.

### 2.5 FLASH-enabled therapeutic indication discovery across diverse disease pathologies

To evaluate the translational utility and cross-domain generalizability of our graph foundation model, we performed a systematic drug repurposing screen across four high-burden diseases with distinct etiologies: Alzheimer’s disease (AD), Parkinson’s disease (PD), Schizophrenia, and Chronic Obstructive Pulmonary Disease (COPD). By spanning neurodegenerative, psychiatric, and respiratory pathologies, these cases serve as a rigorous assessment of the ability of the model to navigate diverse therapeutic landscapes.

We screened 2,869 Food and Drug Administration (FDA)-approved drugs in an inductive setting, utilizing a frozen-encoder linear probing pipeline with a three-class MLP classifier trained on frozen FLASH embeddings(Methods). For each drug–target-disease pair, the model outputs probabilistic scores for each of three classes: therapeutic indication, adverse contraindication, and no association. Using a confidence threshold (*P*_indication_ *>* 0.5), we prioritized candidate drug–disease pairs as high-confidence therapeutic indications for each of the four disease pathologies **(Figs. 5a-c)**.

**Fig. 5.**
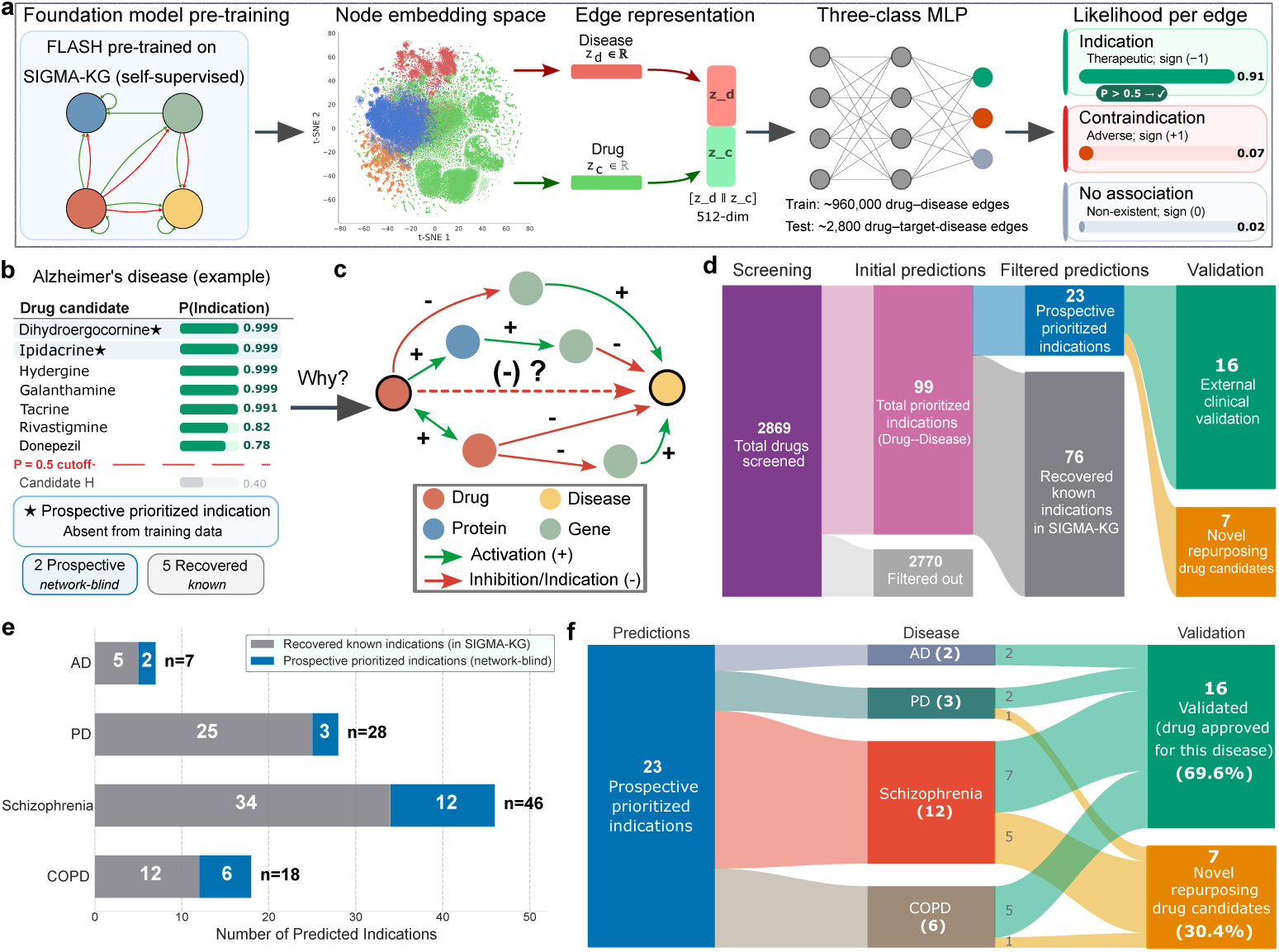
Framework and results of drug repurposing predictions generated by pre-trained FLASH on SIGMA-KG. **a**, Prediction framework. FLASH is first pre-trained self-supervisedly on SIGMA-KG to learn node embeddings. Drug–disease edge representations are then constructed by concatenating the corresponding node embeddings and passed to an MLP to predict the likelihood of indication, contraindication, and no association. **b**, Example predictions for Alzheimer’s disease. The model ranks candidate drugs by predicted probability of therapeutic indication, identifying both approved treatments and high-confidence candidates absent from the training data. **c**, Mechanistic interpretation. Multi-hop biological pathways linking drugs to diseases through proteins and genes are traced within SIGMA-KG, providing mechanistic hypotheses for predicted therapeutic effects. **d**,Repurposing screening workflow. A total of 2,869 drugs were screened across 4 diseases (Alzheimer’s disease (AD), Parkinson’s disease (PD), schizophrenia, and chronic obstructive pulmonary disease (COPD)), yielding 99 initial predicted drug–disease associations. After filtering documented indications present in SIGMA-KG, 23 prospective prioritized indications remained for further evaluation. **e**, Distribution of predicted therapeutics indications across diseases. Predictions include both recovered known indications within SIGMA-KG and prospective network-blind candidates not observed during training. **f**, External validation of prioritized candidates. Among the 23 prospective prioritized indications, 16 correspond to drugs already approved for the predicted disease, providing external clinical support (69.6%), while the remaining 7 represent novel repurposing candidates.

This screening recovered documented clinical knowledge while also identifying prospective therapeutic opportunities **(Fig. 5d)**. In total, FLASH prioritized 99 high-confidence therapeutic indications, providing a map of candidate treatments across all four diseases. To assess the fidelity of our foundation model, we partitioned the 99 prioritized therapeutic indications into recovered and prospective sets based on their presence in the pre-training atlas. We found that FLASH recovered 76 known indications, that is, previously documented drug–disease edges in SIGMA-KG; this indicates that the model preserved and prioritized established pharmacological knowl-edge during embedding learning and downstream classification **(Fig. 5e**, **Table 2 and Supplementary Table 13)**.

**Table 2.**
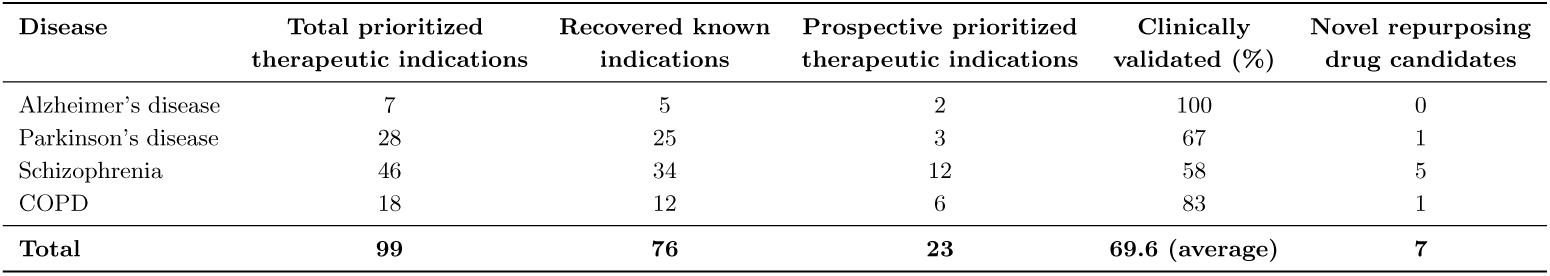
Systematic screening and clinical validation of therapeutic indications across four high-burden diseases. For each disease, 2,869 FDA-approved drugs in the SIGMA-KG universe were screened using pre-trained FLASH embeddings and a three-class MLP classifier. Total prioritized therapeutic indications denote drug–disease pairs with an indication probability *P >* 0.5; Recovered known indications represent predicted edges already documented within the SIGMA-KG training set; Prospective prioritized therapeutic indications correspond to *network-blind* candidates entirely absent from the knowledge graph during pre-training; Clinically validated indicates the subset of prospective candidates retrospectively confirmed via global regulatory records (FDA, European Medicines Agency, Medicines and Healthcare products Regulatory Agency); Novel repurposing drug candidates represent the high-confidence therapeutic hypotheses (including two candidates in Phase 4 trials) prioritized for experimental follow-up. COPD, chronic obstructive pulmonary disease.

| Disease | Total prioritized therapeutic indications | Recovered known indications | Prospective prioritized therapeutic indications | Clinically validated (%) | Novel repurposing drug candidates |
| --- | --- | --- | --- | --- | --- |
| Alzheimer’s disease | 7 | 5 | 2 | 100 | 0 |
| Parkinson’s disease | 28 | 25 | 3 | 67 | 1 |
| Schizophrenia | 46 | 34 | 12 | 58 | 5 |
| COPD | 18 | 12 | 6 | 83 | 1 |
| **Total** | **99** | **76** | **23** | **69.6 (average)** | **7** |

To evaluate the predictive power of FLASH on associations absent from its train-ing data, we then isolated 23 prospective prioritized therapeutic indications (2 for AD, 3 for PD, 12 for Schizophrenia, and 6 for COPD; **Supplementary Table 14**).

These *network-blind* out-of-KG predictions represent drug–disease pairs that were not encoded as indication edges within SIGMA-KG.

Retrospective external clinical validation against global regulatory approval records revealed that the majority (69.6%) of these network-blind predictions correspond to established clinical approvals **(Fig. 5f)**. By systematically searching regulatory databases and clinical registries (e.g., FDA, European Medicines Agency, Medicines and Healthcare products Regulatory Agency; Methods), we found that 16 of the 23 prospective prioritized indications (69.6%) have already received regulatory approval for their respective diseases **(Supplementary Table 14)**. For instance, FLASH correctly identified the acetylcholinesterase inhibitor ipidacrine for AD and the bronchodilator formoterol for COPD, despite these associations being missing from the initial knowledge graph. The high concordance between FLASH’s network-blind pre-dictions and independent regulatory approvals demonstrates the model’s capacity for high-fidelity mechanistic discovery, validating it as a robust framework for bridging systems-level computational insights with real-world therapeutic applications.

The remaining 7 novel repurposing drug candidates represent high-priority ther-apeutic hypotheses generated by the model. Notably, two of these novel candidates for schizophrenia are currently under active Phase 4 clinical investigation for the disease (NCT02593734 and NCT00485823; **Supplementary Table 15**). These findings underscore the utility of sign-aware foundation models in surfacing non-obvious therapeutic opportunities that are both mechanistically grounded and clinically plausible, providing a scalable pipeline for accelerating drug repurposing.

To verify that these candidates arise from biologically plausible mechanisms rather than opaque heuristics, we evaluated the mechanistic interpretability of FLASH by tracing five distinct multi-hop signed metapaths within SIGMA-KG. This approach identifies the gene and protein intermediaries that constitute the signaling cascades for each novel drug–disease indication, providing a transparent molecular basis for the model’s predictions (**Fig. 5c**; Methods). These signed metapaths are intrinsically designed to satisfy structural balance principles, ensuring that the inferred regulatory cascades are mechanistically consistent with biological logic **(Supplementary Fig. 1d)**. For example, FLASH identified a plausible path for COPD (Drug → Protein → Gene → Disease) in which formoterol downregulates the beta-3 adrenergic receptor (encoded by ADRB3), a gene specifically upregulated in the COPD disease state.

By aggregating these intermediate nodes, we identified disease-specific mechanistic target modules that exhibit high biological and functional coherence (a full list of the targets is provided in **Supplementary Table 16**). Comprehensive pathway and protein-protein interaction (PPI) enrichment analyses further confirmed that these target modules are significantly associated with established disease pathophysiology (**Supplementary Fig. 5a-h**; details provided in the Supplementary Information). Together, these results show that our graph foundation model provides not only high-confidence therapeutic prioritization but also a plausible and traceable biological rationale for each drug candidate.

## 3 Discussion

The transition from associative link prediction to sign-aware mechanistic modeling represents a critical shift in systems pharmacology. While traditional graph frameworks successfully capture topological proximities, they frequently fail to resolve the functional polarity (activation/indication versus inhibition/contraindication) that governs biological regulatory systems. Our results show that sign-awareness within FLASH foundation model is essential for accurately resolving the sign composition (transitive logic) that defines multi-step biological signaling. This theoretical advantage is empirically supported by 31 evaluation metrics across three key downstream tasks. Notably, statistical comparisons reveal that the top-performing signed architecture significantly outperformed unsigned models in 27 of the 31 metric evaluations and surpassed relational baselines in 19 instances, highlighting the robust and consistent superiority of sign-aware modeling. By embedding sign-awareness at the representation stage, we mitigate the *polarity drift* inherent in unsigned or purely relational pre-training, ensuring that directional semantics are preserved across diverse biological scales

A core strength of this framework lies in the alignment between Heider’s structural balance theory and biological regulatory motifs. This principle was empirically validated through a systematic drug repurposing screen across four complex diseases: AD, PD, schizophrenia, and COPD. Utilizing a frozen-encoder linear probing protocol, FLASH prioritized 23 network-blind therapeutic candidatesn (Drug–Disease pairs) that were entirely absent from the SIGMA-KG pre-training atlas. Retrospective cross-referencing against global regulatory registries (e.g., FDA and EMA) revealed that 69.6% of these prospective predictions correspond to established clinical approvals, such as ipidacrine for AD and formoterol for COPD (Methods). The predictive power of the model is further underscored by the identification of two candidates for schizophrenia, which are currently under Phase 4 clinical investigation (NCT02593734 and NCT00485823). These results highlights the capacity of sign-aware reasoning to uncover therapeutic overlaps that are mechanistically grounded yet invisible to unsigned methods.

The architectural efficiency of FLASH resolves the historical trade-off between model expressiveness and computational scalability. By utilizing a localized seven-motif aggregation strategy rather than exhaustive higher-order enumeration, FLASH approximates complex relational dependencies with linear scalability. This algorithmic innovation allowed for pre-training on a 3.8-million-edge atlas without the memory-exhaustion failures observed for the signed model SNEA in our largest configurations (Supplemental Information). This establishes FLASH as a scalable transferable foundation model, capable of performing inductive inference on out-of-distribution chemical entities (cold-start DDI prediction; Results) without the need for the task-specific fine-tuning required by predecessors of unsigned GNNs such as TxGNN [5] and signed GNNs such as SIGDR [20].

Beyond predictive accuracy, our framework enables a transparent-box approach to drug discovery. By tracing the algebraic product of signs along multi-hop regulatory paths within SIGMA-KG, FLASH provides testable mechanistic rationales that link molecular perturbations to systemic clinical outcomes. We identified disease-specific target modules that act as molecular intermediaries, linking 7 novel drug candidates to their respective clinical phenotypes. These modules exhibit significant functional inter-connectedness and pathway enrichment which confirms that our mechanistic rationales align with established pathophysiology rather than random topological associations. Such transparency is directly applicable to rational polypharmacy, where signed paths can identify drug combinations that act synergistically on a therapeutic target while neutralizing potential antagonistic effects.

Despite these advances, several limitations remain. The signed edges in SIGMA-KG represent consensus-based, context-averaged effects and may not capture the dynamic sign-reversals seen across different tissues, doses, or disease stages. Furthermore, the reliance on cell-line-derived transcriptomic data introduces a degree of ascertainment bias that may limit direct translation to in vivo human physiology. Additionally, the negative sampling strategy used in three-class benchmarking (Results), while used as standard practice [25], assumes that unobserved associations are true negatives, an assumption that potentially introduces label noise given the inherent incompleteness of current pharmacological databases. Finally, while SIGMA-KG significantly expands chemical coverage, the utility of the model remains constrained by the availability of high-quality transcriptomic signatures for novel chemical series.

Future efforts should prioritize the experimental validation of the novel candidates identified here, such as vigabatrin for PD **(Supplementary Table 13)**, to bridge the gap between computational priority and clinical application. Expanding the SIGMA-KG atlas to include mutations (DNA variants), cell-types and tissues as biomedical entities, and also to incorporate single-cell annd bulk transcriptomics will be essential for moving toward cell-type-resolved and patient-stratified discovery. Ultimately, SIGMA-KG and FLASH provide a scalable, sign-aware foundation for mechanistically informed drug discovery, and we anticipate that the open release of these resources will facilitate more robust predictive modeling in systems pharmacology.

## 4 Methods

### 4.1 Signed multi-omics KG curation

SIGMA-KG was constructed by harmonizing and integrating data from 16 curated biomedical resources **(Supplementary Table 1)** into a unified signed directed heterogeneous KG. It was curated to provide a unified substrate for sign-aware representation learning across molecular and clinical layers. SIGMA-KG comprises 127,422 nodes spanning four biomedical entity types: compounds (76,009 nodes; 59.7%), genes (19,245 nodes; 15.1%), proteins (18,609 nodes; 14.6%), and Diseases/Clinical Phenotypes (13,559 nodes; 10.6%). The KG contains 3,887,829 edges representing twelve distinct relation types organized across inter-layer and intra-layer connections **(Table 1, Supplementary Table 2)**. Critically, each edge carries a signed label (+1/−1) reflecting the polarity of the underlying biological effect. **Fig. 1a** and **Supplementary Fig. 1** present the SIGMA-KG schema and summary statisics.

#### Identifier harmonization

Entity identifiers were harmonized across resources using standard ontologies: HGNC symbols for genes, UniProt accession numbers for proteins, PubChem Compound Identifiers (CIDs) for compounds, and Human Phenotype Ontology (HPO) for diseases/clinical phenotypes. Disease, phenotype and side-effect terms were consolidated using SapBERT biomedical LLM embeddings [27]; term pairs with cosine similarity exceeding 0.8 were merged, reducing 69,207 initial terms to 13,559 unified disease nodes **(Supplemental Information)**.

#### Data sources and integration

Briefly, gene–disease transcriptional dysregulation edges were derived from AMP-AD [28], DiSignAtlas [29], GEO differential gene signatures [30], and Hetionet [6]; compound–gene perturbation edges were derived from LINCS L1000 Connectivity Map consensus signatures [31]; protein-protein interactions were obtained from STRING [32]; compound–disease indication/contraindication edges were integrated from PrimeKG [7], SIDER [33], OnSIDES [34], and repoDB [35]; disease–disease similarity edges were taken from PrimeKG [7]; gene–protein encoding edges were taken from UniProtKB [36]; and compound–protein activation/inhibition edges were taken from IUPHAR [37] and STITCH [38]. Compound–compound similarity edges were derived from the Chemical Checker [39] structural similarity level (A1), which harmonizes data from primary repositories including ChEMBL and PubChem. Complete preprocessing steps and filtering criteria are provided in **Supplementary Information**.

#### Edge construction and sign assignment

Each directed edge was represented as a quadruple (source entity, relation type, sign, target entity), where the sign (+1 or −1) encoded the polarity of the underlying biological or clinical effect. Positive signs were assigned to: *upregulated-in* (gene→disease), *upregulates* (compound→gene), *similar-to* (compound↔compound), *interacts-with* (protein↔protein), *resembles* (disease↔disease), *encodes* (gene→protein), *activates* (compound→protein), and *contraindication* (compound→disease). Negative signs were assigned to: *downregulated-in* (gene→disease), *downregulates* (compound→gene), *inhibits* (compound→protein), and *indication* (compound→disease). This convention encodes contraindications (adverse associations) as positive and indications (therapeutic benefit) as negative, preserving semantics consistent with structural balance theory **(Supplementary Fig. 1d**). For instance, if a compound upregulates (+1) a gene that is downregulated (−1) in a disease, the inferred compound–disease relationship would be negative (−1), consistent with a therapeutic indication.

#### Signed network balance semantics

To ensure the mechanistic and semantic consistency of the integrated multi-omics atlas, we enforced the principles of Heider’s structural balance theory throughout the SIGMA-KG framework. Following identifier harmonization and sign-aware edge construction, we performed a systematic topological refinement to resolve contradictory regulatory paths. Within this signed directed graph, a cycle is defined as structurally balanced if the product of its constituent edge signs is positive:

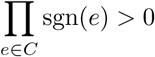

where *C* represents the set of edges in a cycle (e.g., a ”double negative” resulting in net activation). Conversely, cycles yielding a negative product, representing ”frustrated” or biologically inconsistent relations, were resolved through a targeted pruning strategy. For every identified unbalanced cycle, we evaluated the constituent edges based on their source-specific confidence scores (e.g., z-scores from differential expression or curation evidence levels). We preferentially pruned the edge with the lowest confidence score. This pruning was primarily targeted at gene–disease or compound–gene associations, which are inherently noisier than high-confidence compound–disease or gene–protein encoding relations. This optimization step ensures that the resulting atlas provides a consistent substrate for sign-aware graph representation learning, mitigating the propagation of contradictory signals during model pre-training.

### 4.2 FLASH model architecture and pre-training strategy

To enable scalable representation learning over SIGMA-KG, we developed FLASH, a fast and lightweight signed heterogeneous GNN architecture designed to balance expressive power and computational efficiency.

Unlike conventional unsigned or relational GNNs, FLASH explicitly models edge polarity during message passing. The architecture is inspired by Signed Graph Attention Networks (SiGAT) [40], which extend attention mechanisms to signed directed graphs by distinguishing positive and negative neighborhoods. However, directly applying SiGAT to SIGMA-KG is computationally expensive due to the atlas-scale size of the network. We therefore redesign the aggregation strategy to suit the structural and biological properties of SIGMA-KG.

#### Signed neighborhood modeling for SIGMA-KG

While SiGAT exhaustively enumerates dozens of signed motifs to capture higher-order neighborhood patterns of each node, such motif modeling scales poorly for large heterogeneous biomedical graphs like SIGMA-KG. To address this challenge, FLASH defines biologically meaningful motifs in the first-order neighborhood, chosen to preserve signed semantics while ensuring scalability: positive/negative neighbors (2 motifs), positive incoming/out-going neighbors (2 motifs), negative incoming/outgoing neighbors (2 motifs), and positive bidirectional neighbors (1 motif).

The first two motifs capture polarity-specific aggregation without direction. The next four motifs encode polarity together with directionality, which is essential for mechanistic explainability in biological signaling chains (for example, Compound → Gene → Disease). The final motif captures edges such as similarity relations that are semantically positive and have bi-directionality (e.g. protein–protein and compound–compound edges). This seven-motif design preserves the essential signed and directional semantics required for structural balance reasoning while substantially reducing computational complexity compared to exhaustive motif enumeration. This curated seven-motif set serves as the minimal sufficient basis for the model to learn structural balance principles, enabling it to resolve global consistency across the signed multi-omics landscape.

#### Motif-aware attention-based aggregation

Let *G* = (*V, E*) denote SIGMA-KG. For each node *u* ∈ *V*, we initialize its feature representation as **X**(*u*).

Let M denote the motif set, where |M| = 7. For the *i*-th motif *m_i_* ∈ M, we define a motif-aware neighborhood of node *u*:

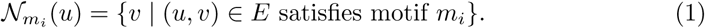

We use the attention-based aggregator in GAT [41] to compute the motif-aware attention coefficient 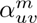 between node *u* and node *v* in N*_m_*(*u*):

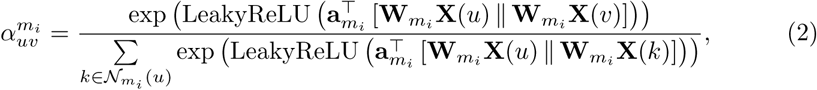

where **a***_mi_* is a weighted vector, **W***_mi_* is a weighted matrix.

The motif-aware attention-based aggregation is then defined as:

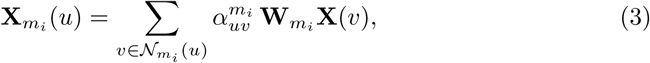

#### Obtain final node embedding

After computing motif-aware representations **X***_m_*(*u*) for all *m* ∈ M, we concatenate them with the original node feature:

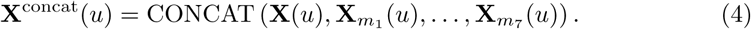

The final node embedding is obtained through a two-layer feedforward network:

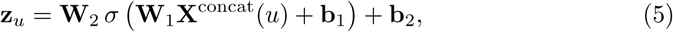

where **W**_1_, **W**_2_ are learnable weight matrices, **b**_1_, **b**_2_ are bias terms, and *σ* is a sigmoid activation function.

#### Self-supervised pre-training objective

FLASH is trained using a self-supervised objective, which models positive and negative relations via a signed proximity objective on the global topology of SIGMA-KG. Specifically, the model optimizes node embeddings by contrasting positive and negative neighborhoods.

For a node *u*, let N (*u*)^+^ and N (*u*)^−^ denote its positive and negative neighbors, respectively. The objective encourages embeddings of positively connected nodes to be similar, while pushing apart those connected by negative edges. The training objective L is defined as:

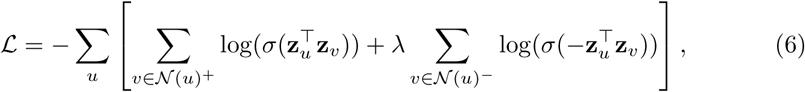

where *σ*(·) is the sigmoid function, and *λ* is a balancing parameter to account for the imbalance between positive and negative edges.

This pre-training strategy explicitly enforces that positively connected nodes have similar embeddings while negatively connected nodes are dissimilar, thereby capturing signed structural patterns in the graph. As a result, the learned embeddings encode multi-hop sign propagation and structural balance properties through signed neighborhood aggregation. Importantly, no task-specific supervision (such as drug–disease labels) is used during pre-training, enabling FLASH to serve as a general-purpose signed graph foundation model transferable across molecular, clinical, and systems-level tasks.

### 4.3 Baselines and fair comparison protocol

To contextualize the performance of FLASH, we benchmarked it against nine GNN methods spanning three architectural categories: (1) unsigned (homogeneous) models (BGRL [42], DGI [43], InfoGraph [44], GraphSAGE [45]); (2) relational heterogeneous models (HGATE [46], DMGI [47]); and (3) signed graph representation learning models (SGCN [48], SNEA [49], SDGNN [50]). All baselines were implemented within a self-supervised pre-training framework to generate task-agnostic latent node representations for downstream evaluation. To ensure a fair and optimized comparison, each model was trained using the specific hyperparameter configurations recommended by their original authors (**Supplementary Table 17**).

To ensure a fair and rigorous comparison, all models were initialized with identical 256-dimensional node features derived from a truncated SVD of the adjacency matrix. We standardized the output at 256-dimensional node embeddings across all baselines; this dimensionality was selected as a necessary computational ceiling, as several complex architectures (specifically SNEA, SDGNN, and HGATE) consistently encountered memory-exhaustion failures on the 3.8M-edge SIGMA-KG atlas when attempting to generate 512-dimensional embeddings.

The information accessible to each model was strictly controlled to isolate the impact of signed multi-relational structures. Unsigned baselines were trained on an unsigned version of SIGMA-KG with all edge types and polarities discarded. Relational baselines utilized the full 12-relation multiplex structure, while signed baselines and FLASH received the signed network, with edges labeled as +1 or −1.

Model evaluation was conducted using a dual-stratified protocol across three independent replicates with matched random seeds to ensure statistical reproducibility. For molecular (Ts-MoA) and clinical (DCR) tasks, we employed an internal held-out validation strategy. Specifically, SIGMA-KG was partitioned into a training subgraph and an independent set of evaluation edges (compound–protein or compound–disease). Each foundation model was first pre-trained in a self-supervised manner on the training subgraph to generate node embeddings . Subsequently, an MLP classifier (binary and three-class) was trained on the training edges using these frozen embeddings and then evaluated on the held-out test set.

For systems-level DDI prediction, we implemented a more stringent external validation protocol. Baseline models were pre-trained on the complete SIGMA-KG atlas, and the resulting compound embeddings were directly transferred to two independent, unseen DDI benchmark datasets: DrugBank (version 5.1) [23] and MUDI [24]. To eliminate downstream classifier variance as a confounding factor, we employed a standardized MLP architecture and identical hyperparameters for FLASH and all baseline counterparts across every task **(Supplementary Table 18)**. By standardizing the classifier and using frozen embeddings, we ensure that observed performance deltas are strictly attributable to the architectural inductive biases of the foundation models rather than variance in downstream supervised learning.

### 4.4 Downstream evaluation for edge-sign prediction via frozen-encoder linear probing

To rigorously assess the predictive utility of the FLASH foundation model, we implemented a standardized benchmarking framework across two critical pharmacological scales: molecular-level Ts-MoA and clinical-level DCR. Both tasks were formulated as edge-sign prediction problems using a frozen-encoder linear probing protocol. This design isolates the quality of the learned representations from the complexity of the downstream classifier, ensuring that performance gains are directly attributable to the sign-aware inductive biases of the foundation model.

#### 4.4.1 Unified benchmarking protocol

For both the Ts-MoA and DCR tasks, we followed a consistent stratified evaluation pipeline to ensure that the foundation model remained entirely naive to evaluation data during pre-training:

- **Task-specific graph construction and data leakage prevention:** Prior to self-supervised pre-training, all labeled edges for the target task (e.g., Compound–Protein or Compound–Disease) were randomly partitioned into training, validation, and test sets using a stratified split (0.8*/*0.1*/*0.1). To prevent information leakage, a task-specific pre-training KG was constructed by starting from the full SIGMA-KG and removing the validation and test edges corresponding to the specific task. The training edges and all remaining relation types in sigma-kg were retained. Initial node features (truncated SVD) were computed exclusively from this restricted pre-training topology.
- **Linear probing architecture:** Following pre-training, node embeddings were extracted and frozen. For any candidate edge (*u, v*), a 512-dimensional feature vector was generated by concatenating the embeddings of the source and target nodes [**z***_u_* ∥ |**z***_v_*]. This vector served as the input for a standardized MLP head. The classifier was optimized using Binary Cross-Entropy (BCE) loss for binary classification tasks and Categorical Cross-Entropy for three-class evaluations. For each task, the MLP was trained on the training edge split, with early stopping based on validation-set performance (Macro-F1). Final performance was measured exclusively on the 10% held-out test set.
- **Three-class open-world evaluation:** To simulate realistic discovery settings, we introduced a ”non-existent” class (0) by uniformly sampling node pairs (compound–protein or compound–disease) absent from SIGMA-KG. Following established protocols [25], we maintained a 5:1 ratio of non-existent edges to signed edges. The MLP was thus trained to distinguish between three classes: activation/contraindication (+1), inhibition/indication (−1), and non-existent (0).
- **Replication and evaluation:** To ensure statistical rigor, the entire pipeline, including graph partitioning, pre-training, and MLP evaluation, was repeated over three independent runs (matched random seeds for stratified edge split). Final metrics are reported as the mean ± standard deviation of AUROC, AUPRC, Accuracy, and Macro-F1 scores across these runs.

#### 4.4.2 Ts-MoA prediction

The Ts-MoA task evaluates the model’s ability to distinguish the functional polarity of drug–target engagement. We labeled compound–protein edges based on their functional effect, where activation/agonist corresponds to a positive sign (+1) and inhibition/antagonist corresponds to a negative sign (−1). The dataset comprised 13,711 high-confidence interactions (6,389 activation and 7,322 inhibition), providing a balanced distribution for evaluating mechanistic reasoning. Binary and three-class classification results are detailed in **Supplementary Table 5** and **Supplementary Table 6**, respectively.

#### 4.4.3 DCR prediction

At the clinical scale, we evaluated the model’s capacity to bridge molecular perturbations with clinical phenotypes. Consistent with the signed semantics of SIGMA-KG, compound–disease edges were labeled as contraindication (+1; adverse association) or indication (−1; therapeutic association). The DCR dataset represents a severely imbalanced setting, containing 11,896 indication and 148,061 contraindication edges (approx. 1 : 12.4 ratio). To address this, the MLP loss function was class-weighted inversely to the frequencies observed in the training split. Results are reported in**Supplementary Table 7** and **Supplementary Table 8**.

### 4.5 Drug-drug interaction prediction on external resources

We evaluated whether pre-trained embeddings generalize to DDIs absent from SIGMA-KG, using two external benchmark datasets whose interaction pairs were entirely unseen during pre-training.

#### DDI benchmark datasets

Two benchmark datasets were used: DrugBank (version 5.1) [23] and MUDI [24]. The DrugBank dataset, manually curated from FDA and Health Canada drug labels and primary literature, annotates pairwise interactions across 113 relation types capturing metabolic, therapeutic, and adverse effect relationships. The MUDI dataset, constructed from a pharmacodynamic perspective using pharmacological text and multimodal chemical representations, annotates interactions as one of three clinically meaningful categories: *Synergism*, *Antagonism*, or *New Effect*. The unprocessed DrugBank dataset contained 222,696 interacting pairs across 1,876 compounds; the unprocessed MUDI dataset contained 310,532 interacting pairs across 1,198 compounds.

#### Pre-processing

Compound identifiers were harmonized to PubChem CIDs via the PubChem Identifier Exchange Service (Supplementary Methods). Self-loop edges (a compound paired with itself) were removed, and pairs with multiple interaction-type annotations were deduplicated to a single binary interaction label. To prevent inverse-pair leakage across data splits, edges were treated as unordered: each pair was canonicalized as (min{*c_i_, c_j_*}, max{*c_i_, c_j_*}) by sorting compound identifiers. Subsequently, only drug pairs for which both compounds were present in SIGMA-KG were retained. After preprocessing, DrugBank dataset comprised 195,967 interacting pairs across 1,645 compounds; MUDI dataset comprised 274,510 pairs across 1,129 compounds. For MUDI, the original train/test split was discarded in favor of the unified protocol below.

#### Task formulation

Although both benchmark datasets provide multi-class interaction annotations, we formulated DDI prediction as a binary edge classification task (a pair of drugs interact vs. does not interact) for three reasons: (i) to maintain a consistent evaluation protocol across resources with different and non-aligned label taxonomies, and (ii) because the primary objective was to assess whether pre-trained embeddings capture general pharmacological relatedness rather than to discriminate among specific interaction mechanisms and (iii) both benchmarks exhibit severe class imbalance: in DrugBank, the three most prevalent interaction types comprise 66% of all pairs, whereas the majority of categories each constitute less than 1%; in MUDI, the *Synergism* type alone accounts for 83% of all annotations. This extreme distribution skew renders multi-class evaluation statistically unreliable for minority classes, justifying our focus on binary classification protocols.

#### Embedding extraction and negative sampling

Each baseline was trained on the full SIGMA-KG in a self-supervised manner to obtain 256-dimensional node embeddings, which remained frozen during downstream evaluation. For each external dataset, only embeddings for drugs present in that preprocessed dataset were used. Positive edges corresponded to annotated interacting drug pairs from each dataset; negative edges were generated by randomly sampling unordered drug pairs– *from the set of drugs present in each preprocessed dataset* –that were not annotated as interacting, at a 1:1 ratio relative to positive pairs. This negative-sampling strategy implicitly assumes that unannotated pairs are non-interacting–a standard but imperfect assumption that may introduce label noise, particularly for incompletely curated resources.

#### Transductive (warm-start; random pair split) evaluation scenario

For each dataset and each random seed, the combined positive and negative edge set was randomly partitioned into training/validation/test splits (0.8/0.1/0.1, stratified by class). As splits are edge-wise random, drugs may appear in both training and test sets (transductive setting). Each candidate pair (*c_i_, c_j_*) was represented by concatenating embeddings in canonical order, yielding a 512-dimensional feature vector [**z**_min(*c*_*_i,cj_* _)_ ∥| **z**_max(*c*_*_i,cj_* _)_]. A binary MLP classifier (minimizing binary cross-entropy, with early stopping on validation Macro-F1) was trained to predict interaction status (interacting versus non-interacting); architecture and hyperparameters were held constant across baselines (**Supplementary Table 18)**.

#### Inductive (cold-start) evaluation scenario

To assess generalization to unseen drugs, we implemented a single-endpoint cold-start protocol. For each dataset and each random seed, the unique drug set *D* was partitioned into disjoint subsets *D*_train_, *D*_val_, and *D*_test_ (0.8/0.1/0.1). Training positives comprised interactions between drugs both in *D*_train_. Validation positives comprised interactions in which at least one endpoint belonged to *D*_val_ and the other to *D*_train_ ∪ *D*_val_. Test positives comprised interactions in which at least one endpoint belonged to *D*_test_ and the other to *D*_train_ ∪ *D*_test_.

For each split, negatives pairs were sampled from the corresponding drug universe at a 1:1 ratio, subject to the same unannotated-equals-negative assumption. This design ensures that every validation and test edge involves at least one drug absent from classifier training, approximating the practical scenario of predicting interactions for a newly introduced compound. MLP architecture and hyperparameters were held constant across baselines (**Supplementary Table 18**).

#### Replication and evaluation

For each baseline and dataset, data partitioning, negative sampling, and MLP training/evaluation were repeated across three independent random seeds; mean ± standard deviation are reported. The same seeds were used across baselines to ensure matched comparisons. For a given setting, the MLP architecture and hyperparameters were held constant across all baselines **(Supplementary Table 4)**. Evaluation metrics included AUROC, AUPRC, accuracy, macro-F1, and class-specific F1. Full results for the transductive and inductive evaluation scenarios are reported in **Supplementary Tables 9and 10**, and **Supplementary Tables 11and 12**, respectively.

### 4.6 Computational efficiency and scalability benchmarking

To evaluate the computational efficiency and scalability of FLASH, we benchmarked its training time and peak GPU memory usage against nine GNN baselines spanning three architectural categories. All experiments report training time and peak GPU memory consumption measured over 100 epochs **(Supplemental Information; Supplementary Fig. 4)**.

#### Subnetwork construction

Scalability was assessed on subnetworks sampled from SIGMA-KG with systematically varied sizes. For node-scaling experiments, we varied the number of nodes from 20,000 to 120,000 while maintaining a constant average degree of 30.51 (matching the global average degree of SIGMA-KG). For edge-scaling experiments, we fixed the number of nodes at 100,000 and varied the number of edges from 500,000 to 2,500,000. This design isolates the impact of graph order (node count) and graph size (edge count) on computational cost.

#### Training protocol

All models were trained in a self-supervised manner on each subnetwork using identical initial node features (128-dimensional truncated SVD of the adjacency matrix), the same number of epochs (100), and produced 128-dimensional node embeddings. Homogeneous baselines were trained on unsigned versions of the subnetworks in which edge signs and relation types were discarded.

#### Hardware environment

All experiments were conducted on a Linux workstation equipped with dual AMD EPYC 7453 28-core processors (112 logical cores, up to 3.49 GHz), 2 TiB of RAM, and an NVIDIA L40S GPU (48 GB). Each training run was executed on a single dedicated GPU with no concurrent jobs to ensure consistent timing measurements.

### 4.7 Drug repurposing

#### 4.7.1 Case studies

##### Disease selection

To evaluate the translational utility of the foundation model, we applied FLASH to identify novel therapeutic indications for five diseases representing diverse pathophysiological mechanisms: AD (neurodegenerative), PD (neurodegenerative), schizophrenia (psychiatric), COPD (respiratory). This selection was designed to assess generalization across distinct therapeutic areas and molecular etiologies.

##### Drug candidate universe

To focus the analysis on drugs with prior regulatory approval, the candidate drug universe was restricted to the 2,869 compounds in SIGMA-KG with at least one established clinical association. We systematically screened this universe against each target disease to prioritize candidate drugs for novel therapeutic indications.”

##### Edge representation and MLP classifier setup

Drug repurposing was formulated as a three-class edge classification task **(5a)**. We utilized 256-dimensional FLASH node embeddings, pre-trained on the full SIGMA-KG in a self-supervised manner, as frozen input features. For any given drug-disease pair, an edge representation was constructed by concatenating the drug embedding **z***_c_* and the disease embed-ding **z***_d_* into a 512-dimensional feature vector [**z***_c_*∥|**z***_d_*]. These vectors were fed into a MLP with a softmax output layer to predict three classes: (i) indication (therapeutic association; label −1), (ii) contraindication (adverse association; label +1), and (iii) non-existent (no known association; label 0).

##### Training strategy and class imbalance mitigation

We implemented a stratified five-fold CV strategy. The training and validation sets were constructed from the 159,957 existing drug-disease edges in SIGMA-KG (labeled as indication or contraindication). To represent the non-existent class, we performed negative sampling by randomly pairing drug nodes (from the pool of 2,869) and disease nodes (from the pool of 13,559) that lacked any established association in SIGMA-KG. The number of sampled non-existent edges was set to five times the number of labeled edges; these were then uniformly distributed across the five folds. Notably, the ratio of indication to contraindication edges in SIGMA-KG is approximately 1:12.4. To account for this substantial class imbalance, the MLP was trained using a weighted cross-entropy loss function, with class weights set inversely proportional to their respective frequencies in the training data. MLP architecture and hyperparameter details are provided in Supplementary Methods.

##### Disease-specific screening and test set construction

For each of the five target diseases, a separate screening pipeline was executed. While the model architecture and the global training/validation folds remained consistent across all five pipelines to ensure a generalized understanding of pharmacological patterns, the test set differed according to the target disease. Specifically, the test set for a given disease (e.g., AD) consisted of all 2,869 candidate drug–disease edges formed by pairing that disease node with every drug in the candidate universe—yielding edges such as AD–Drug_1_, AD–Drug_2_, . . ., AD–Drug_2869_. This process was repeated independently for each target disease (AD, PD, Schizophrenia, and COPD).

##### Cross-validation and hyperparameter optimization

During each fold of the CV, the MLP was trained on four folds (with the validation fold used for early stopping based on macro-F1) and then applied to the disease-specific test set, outputting softmax probabilities for each of the three classes for every test edge. Consequently, each testing edge (e.g., AD-Drug *X*) received five separate softmax probability distributions. MLP hyperparameters were tuned via nested five-fold CV on the training folds. Architecture and hyperparameter details are provided in Supplementary Table 18.

##### Prioritized therapeutic indications

We calculated the final confidence score for each candidate indication by averaging the “indication” class probabilities across all five CV folds. Edges with a mean probability *P* (indication) *>* 0.5 were retained as prioritized therapeutic indications.

##### prospective prioritized therapeutic indications

To ensure the discovery of novel repurposing candidates, we applied a rigorous filtering procedure: any predicted indication edge (drug–disease) already present in the SIGMA-KG (recovered known indications) was excluded from the list of prioritized therapeutic indications. Only network-blind predictions, drug–disease pairs absent from the pre-training KG, were retained as *prospective prioritized therapeutic indications*.

##### External clinical validation of prospective prioritized therapeutic indications

Retrospective clinical validation was performed by cross-referencing predicted drug–disease pairs against regulatory approval records from FDA, European Medicines Agency (EMA), Medicines and Healthcare products Regulatory Agency (MHRA), and State Agency of Medicines of the Republic of Latvia (SAM), using standardized drug and disease identifiers. Validation was conducted in a systematic manner. A drug–disease pair was considered validated if the drug is officially approved for the indication in at least one of the referenced databases using standardized identifiers.

#### 4.7.2 Mechanistic explainability and target inference

##### Metapath-based target identification

To interpret the molecular basis of the predictions, we identified intermediate nodes connecting predicted drugs to their respective target diseases through SIGMA-KG. We investigated five specific multi-hop metapaths based on shortest-path connectivity:

1. Drug → Gene → Disease
2. Drug → Protein → Gene → Disease
3. Drug → Drug → Gene → Disease
4. Drug → Drug → Protein → Gene → Disease
5. Drug → Drug → Protein → Protein → Gene → Disease

##### Pathway and PPI enrichment analysis

Functional enrichment analysis was performed on the aggregated gene and protein targets for each disease using the STRING database [32]. Pathway enrichment was assessed with a FDR threshold of *<* 0.0001. PPI enrichment analysis was performed to evaluate the biological coherence of the target sets. Significant PPI enrichment *p*-value indicates that the identified targets form a functionally interconnected module rather than a random collection of proteins, supporting the mechanistic plausibility of the foundation model’s predictions.

### 4.8 Statistical Analysis

To rigorously assess the significance of the performance gains achieved by FLASH, all experiments were executed across three independent replicates (*N* = 3) using distinct random initializations to ensure reproducibility. To maintain a fair comparison, the same three random seeds were applied across all baselines for data partitioning and stochastic sampling. Statistical significance was evaluated using the unpaired Welch’s *t*-test, which is robust to potential differences in variance between model performances.

We implemented a two-tiered comparative analysis to evaluate the results:

1. **Inter-class comparison:** We compared the top-performing signed baseline against the best-performing unsigned and relational baselines using a one-sided unpaired Welch’s *t*-test. This analysis was designed to specifically validate the inductive bias afforded by explicit edge polarity across different tasks (**Supplementary Table 3**).
2. **SOTA validation:** For each metric, we conducted a direct comparison between FLASH and the runner-up baseline (in cases where FLASH established SOTA results) or the top-performing baseline (if FLASH matched existing performance). These comparisons utilized a two-sided unpaired Welch’s *t*-test to establish statistical superiority or parity (**Supplementary Table 4**).

Statistical significance was predefined as *P <* 0.05. Results are reported as mean ± standard deviation (s.d.). Where FLASH significantly outperformed the leading baselines, significance is denoted using a hierarchical notation: ^∗^ for *P <* 0.05, ^∗∗^ for *P <* 0.01, and *n.s.* (not significant) for *P* ≥ 0.05. In instances where differences did not reach the significance threshold, FLASH is described as achieving performance comparable to existing SOTA methods. All statistical computations and distribution analyses were performed using SciPy (v1.12.0) [51]

### Data availability

All datasets utilized in this study are publicly available.

- Multi-omics and perturbation signatures were accessed via the following repositories: AMP-AD studies were obtained through the AD Knowledge Portal (https://adknowledgeportal.org; accession no. syn21241740); DiSignAtlas signa-tures are available at http://www.inbirg.com/disignatlas/; Chemical Perturbations Consensus Signatures (LINCS L1000) and GEO signatures were retrieved from Harmonizome (https://maayanlab.cloud/Harmonizome/); and multilevel biological signatures for chemical compounds across 25 biological levels were retrieved from the Chemical Checker portal (https://chemicalchecker.org).
- Clinical indications and trial outcomes were sourced from repoDB (http://apps.chiragjpgroup.org/repoDB/). Phenotypic indications were sourced from PrimeKG via the Harvard Dataverse (https://doi.org/10.7910/DVN/DC7YDX). Adverse drug reaction data were obtained from SIDER 4.1 (http://sideeffects.embl.de/) and the NLP-extracted OnSIDES database (https://nsides.io/).
- Biological interaction networks and knowledge graphs were integrated from: Hetionet (https://github.com/hetio/hetionet); the IUPHAR/BPS Guide to PHARMA-COLOGY (https://www.guidetopharmacology.org); STITCH 5.0 (http://stitch.embl.de/); STRING v12.0 (https://string-db.org/); and UniProtKB (https://www.uniprot.org/).

## Author contributions

M.M. prepared data, implemented algorithms, performed experiments, analyzed data and co-wrote the manuscript. S.Z. analyzed data and co-wrote the manuscript. I.A. performed experiments. P.Z. analyzed data. L.X. conceived methods, planned experiments and co-wrote the manuscript.

## 1 Related Work

The evolution of biomedical knowledge graphs (KGs) has transitioned from bipartite maps to large-scale multi-relational atlases [1]. While resources such as Hetionet [2], SPOKE [3], and PrimeKG [4] have enabled systemic drug-disease modeling, they remain fundamentally restricted by a ”polarity-blind” architecture. These KGs primarily encode approved compounds (*∼*1,552 in PrimeKG), whereas SIGMA-KG expands this to 76,009 small molecules, providing the necessary chemical breadth for early-stage discovery. Furthermore, existing KGs conflate biological activation and inhibition into unsigned proximities (unsigned edges), a limitation that prevents models from distinguishing therapeutic efficacy from toxicity [5, 6]. While curated databases such as SIGNOR 3.0 [6] and TRRUST v2 [7] encode signed interaction polarities (*±*1) in their data, they are modality-specific and lack the multi-omics integration required for atlas-scale representation learning.

Graph neural networks (GNNs) have emerged as the dominant paradigm for learning over biomedical KGs, enabling powerful representations (node embeddings) for downstream tasks including, drug-target interaction [8], drug–drug interaction [9], genotype-phenotype association predictions [10], and drug repurposing [11]. Homogeneous unsigned methods, including GraphSAGE [12], Infograph [13], and DGI [14], learn representations through self-supervised objectives such as mutual information maximization or bootstrapped consistency, without requiring labeled data. Relational GNNs (rGNNs), such as R-GCN [15], HGT [16], and HGATE [17], treat ”activation” and ”inhibition” as independent categorical labels. This categorical treatment is architecturally incapable of algebraic sign-composition (e.g., *A −→ B −→ C* =*⇒ A −*^+^*→ C*; structural balance theory) [18], leading to the ”polarity drift” identified in our Introduction. In contrast, path-based reasoners like BioPathNet [19] attempt to capture multi-hop logic but rely on supervised task-specific labels, limiting their generalizability across diverse biological contexts.

The field is currently shifting toward graph foundation models. TxGNN [11] represents a major milestone in zero-shot drug repurposing, utilizing metric learning (e.g., RotatE [20]) on unsigned relational graphs. However, TxGNN requires supervised fine-tuning on labeled drug-disease pairs and cannot perform mechanistic reasoning across signed regulatory paths. Similarly, self-supervised models such as PT-KGNN [21] demonstrate scalability on unsigned KGs but apply homogeneous aggregation that discards functional edge polarity. FLASH addresses this by embedding sign-awareness into the self-supervised pre-training objective itself, producing representations that encode causal logic without task-specific supervision.

Existing signed GNN (sGNN) architectures, such as SGCN [22], SNEA [23], and SDGNN [24], were primarily developed for social networks where edge signs reflect ”friend/enemy” sentiments. These models often suffer from high computational complexity when applied to millions of edges and typically utilize transductive or supervised objectives that preclude ”cold-start” discovery for unseen compounds [5]. Recent biological applications such as SIGDR [25] and CSGDN [10] have shown the promise of sign-aware learning for drug-disease and gene-phenotype prediction downstream tasks, yet they operate on single-modality networks of restricted scale.

FLASH represents a necessary evolution: it is, to our knowledge, the first lightweight sGNN architecture designed for heterogeneous multi-omics graphs at an atlas scale (over 3.8M edges), bridging the gap between scalable self-supervised learning and the sign-consistent logic of biological signaling systems.

## 2 Computational efficiency and scalability analysis

To rigorously evaluate the scalability of FLASH, we conducted a comparative bench-marking study against nine GNN baselines spanning unsigned, relational, and signed architectural categories. All experiments were performed under controlled conditions to ensure a fair assessment of training time and peak GPU memory consumption.

### Subnetwork Construction and Scaling Protocol

Scalability was assessed using subnetworks sampled from SIGMA-KG with systematically varied dimensions. For node-scaling experiments, we varied the number of nodes from 20,000 to 120,000 while maintaining a constant average degree of 30.51 to match the global topology of the full atlas. For edge-scaling experiments, the node count was fixed at 100,000 while the number of edges varied from 500,000 to 2,500,000. This dual-axis design allows for the isolation of computational costs associated with graph order (nodes) versus graph size (edges).

### Runtime scaling

FLASH exhibited favorable scaling behavior across both dimensions (Fig. 4a,c). Training time increased approximately linearly with network size, completing in 55 seconds on graphs with 2.5 million edges and 93 seconds on graphs with 120,000 nodes. In contrast, sGNN baselines such as SNEA and SGCN required 531 and 409 seconds, respectively, at equivalent edge counts—representing 7- to 10-fold longer training times. Relational baselines (HGATE, DMGI) showed intermediate performance but with steeper scaling, while lightweight unsigned methods (BGRL, InfoGraph) achieved faster runtime but forgo the signed and relational structure that FLASH exploits.

### Memory scaling

Peak memory consumption followed similar trends: at 2.5 million edges, FLASH required approximately 10.8 GB of GPU memory, compared with 15.6 GB for SNEA, 17.3 GB for SDGNN, 16.3 GB for DMGI, and higher for other signed models (Fig. 4b,d).

### Analysis of results

The results (summarized in Fig. 4) highlight that FLASH avoids the cubic or quadratic complexity often associated with motif-aware or relational attention mechanisms. By utilizing a streamlined seven-motif aggregation strategy, FLASH achieves runtime performance comparable to lightweight unsigned models while outperforming specialized sGNNs in both speed and memory retention. This efficiency is critical for handling the increasing density of multi-omics data.

## 3 Functional enrichment and protein-protein interaction (PPI) analysis of mechanistic target modules

### Mechanistic target modules exhibit high biological coherence and disease-specific relevance

To validate the explainability of the therapeutic indication predictions, we aggregated the intermediate genes and proteins identified through multi-hop metapath tracing (Methods) into disease-specific mechanistic target modules. We performed Gene Set Enrichment Analysis (GSEA) and PPI enrichment analysis for each set of targets to determine if the model’s predictions converged on established disease biology. Table 16 contains the complete list of disease-specific mechanistic target (genes and proteins).

### Pathway analyses support the mechanistic plausibility of prioritized drug candidates

For each disease, the identified mechanistic target modules showed highly significant enrichment in pathways consistent with known pathophysiology (FDR ¡ 0.0001; Fig ??). Alzheimer’s disease targets were predominantly associated with amyloid fibril formation and calcium homeostasis, while Parkinson’s disease targets were enriched in dopaminergic synapse function and amino acid metabolism. Similarly, schizophrenia-associated modules were localized to serotonergic and cholinergic synapse pathways, and COPD targets were enriched in smooth muscle contraction and corticotropin hormone signaling.

### PPI enrichment analysis revealed that the target sets for all four diseases exhibit significantly higher connectivity than expected by chance

(*P <* 1.0 *×* 10*^−^*^16^ for PD, Schizophrenia, and COPD; *P* = 0.003 for AD). The formation of these dense functional modules suggests that FLASH effectively captures the ”system-level” impact of drugs on disease-specific protein networks. These results, detailed in Fig **??**, provide evidence that the FLASH foundation model makes sound, explainable predictions rooted in the modular architecture of human biology.

## 4 SIGMA-KG constructions details

### 4.1 SIGMA-KG Data sources and entity harmonization

SIGMA-KG integrates 16 publicly available biomedical resources (Table 1). All data were downloaded between [November 2024] and [August 2025]. Gene identifiers were mapped to HGNC symbols via HGNC BioMart []. Protein identifiers were standardized to UniProt accessions using the UniProt ID mapping tool (https://www.uniprot.org/id-mapping/). Compound identifiers were harmonized to PubChem Compound Identifiers (CIDs) using the PubChem Identifier Exchange Service (https://pubchem.ncbi.nlm.nih.gov/idexchange/idexchange.cgi). Input identifiers included compound names (synonyms), DrugBank identifiers, SMILES strings, and InChIKeys depending on the source. To ensure a unique canonical identifier per compound and avoid duplication arising from stereoisomers or salt forms, the Parent CID was retrieved for each entry. Compounds lacking a valid PubChem CID mapping were excluded. Disease terms, harmonized to Human Phenotype Ontology (HPO), were consolidated using SapBERT biomedical LLM [26]: each term was encoded as a 768-dimensional embedding, pair- wise cosine similarities were computed, and terms with similarity *≥* 0.8 were merged via single-linkage clustering, reducing 69,207 terms to 13,559 nodes.

### 4.2 Data sources and SIGMA-KG edge construction details

- **Gene–Disease.** Differential expression data were obtained from AMP-AD, DiSig-nAtlas, GEO, and Hetionet. Z-scores were computed from normalized log_2_ fold-change values. Edges with *z >* +2 were assigned *upregulated-in* (+1); edges with *z < −*2 were assigned *downregulated-in* (*−*1). Direction: gene*→*disease.
- **Compound–Gene.** Consensus Perturbation signatures were obtained from LINCS L1000 CMAP Chemical Perturbation Consensus Signatures (12,126 genes and 23,913 small chemical perturbations). The dataset was filtered using gene-wise z-scoring to include associations (edges) with *|z| ≥* 3. Edges with positive *z* were assigned *upregulates* (+1); edges with negative *z* were assigned *downregulates* (*−*1). Direction: compound*→*gene.
- **Compound–Compound.** Structural similarity edges were sourced from the Chemical Checker (CC) [27], a comprehensive resource that categorizes small-molecule bioactivity into five levels. We utilized CC Level 1 (Chemistry) data, specifically the A1 coordinate representing 2D structural fingerprints. The extensive scale of SIGMA-KG (76,009 compounds) results from our integration strategy: we initially considered the full CC universe of *≈*800,000 small molecules and filtered for pairs where at least one compound was present in our multi-omics layers (transcriptomics, proteomics, or clinical). We retained bidirectional edges for compound pairs exhibiting a Tanimoto similarity score *≥* 0.8 based on ECFP4 signatures. Chemical identifiers provided by CC in InChIKey format were mapped to PubChem CIDs using the PubChem Identifier Exchange Service to ensure consistency across the atlas. This process yielded 581,872 high-confidence structural similarity links.
- **Protein–Protein.** Interactions were obtained from STRING with combined confidence *≥* 0.8. Assigned *interacts-with* (+1). Direction: bidirectional.
- **Compound–Disease.** Clinical associations were aggregated from PrimeKG, Side Effect Resource (SIDER) (version 4.1, Frequency Percentage *≥* 0.4), OnSIDES (version 3.1.0), and Drug Repositioning DataBase (repoDB). SIDER (4.1) includes information on marketed drugs and their recorded adverse drug reactions, OnSIDES is a database of adverse drug events extracted from drug labels using natural language processing (NLP), and repoDB contains a standard set of drug repositioning successes and failures extracted from FDA documents. Therapeutic relationships were assigned *indication* (*−*1); adverse relationships were assigned *contraindication* (+1). Direction: compound*→*disease.
- **Gene–Protein.** Encoding relationships were obtained from UniProtKB, restricted to reviewed human entries. Assigned *encodes* (+1). Direction: gene*→*protein.
- **Compound–Protein.** Functional modulation data were obtained from IUPHAR/BPS Guide to Pharmacology database (v2025.3) and STITCH (v5.0, Combined Score *≥* 0.85). Agonist annotations were assigned *activates* (+1); antagonist annotations were assigned *inhibits* (*−*1). Direction: compound*→*protein.
- **Disease–Disease.** Similarity edges were obtained from Phenotype-Phenotype positive associations in PrimeKG extracted from HPO. Assigned *resembles* (+1). Direction: bidirectional.

**Quality control.** All duplicated edges with conflicting signs were removed. Duplicate edges from multiple sources were collapsed. Isolated nodes (node with zero connections) were removed. The final signed KG was exported as directed edge lists in the format: (Source entity, Relation, Sign, Target entity).

### 4.3 Detailed baseline model descriptions and training hyperparameters

To evaluate the performance of FLASH, we benchmarked it against nine state-of-the-art self-supervised graph neural network (GNN) baselines. All models were pre-trained using the hyperparameter configurations and loss objectives recommended in their respective original publications to ensure a robust and fair comparison. Baseline-specific hyperparameters are provided in Table 17.

- **Bootstrapped Graph Latents (BGRL)**: This framework utilizes a student-teacher architecture to predict alternative augmentations of the same input graph. The training objective is a cosine similarity loss, which eliminates the necessity for negative sampling [28].
- **Deep Graph Infomax (DGI)**: This method maximizes mutual information between local node patches and a global graph summary. It employs a contrastive objective using a discriminator to distinguish between real and corrupted graph structures [14].
- **InfoGraph**: This model learns graph-level representations by maximizing mutual information between global embeddings and substructures across multiple scales. The objective ensures the encoding of features shared across nodes, edges, and larger graph components [13].
- **GraphSAGE**: This inductive framework aggregates local neighborhood features to learn functional representations instead of fixed node embeddings. It is optimized via a graph-based loss that encourages embedding proximity for topologically connected nodes [12].
- **Deep Multi-View Graph Infomax (DMGI)**: This approach handles multiplex networks by maximizing mutual information across relation-specific views. It uses consensus regularization and a universal discriminator to minimize disagreements between different relation types [29].
- **Heterogeneous Graph Attention Auto-Encoder (HGATE)**: This architecture reconstructs heterogeneous graph topology and node attributes via a hierarchical attention mechanism. The loss objective consists of dual reconstruction errors for edges and attributes across diverse meta-paths [17].
- **Signed Graph Convolutional Network (SGCN)**: This model generalizes GCNs to signed graphs by separately aggregating information from positive and negative neighbors. It is optimized using a principled loss function that constrains embeddings to satisfy structural balance theory [22].
- **Signed Network Embedding via Attention (SNEA)**: This framework employs a masked self-attention mechanism to weigh the importance of neighbors connected by different link polarities. The objective uses a sign-aware link prediction loss to preserve complex signed network interdependencies [23].
- **Signed Directed Graph Neural Network (SDGNN)**: This model captures link signs, directions, and triad structures by incorporating sociological theories of status and balance. The objective is a multi-task reconstruction loss optimizing for edge sign, directionality, and signed triangle consistency [24].

### 4.4 Classifiers (MLPs) architecture and hyperparameters

To ensure a rigorous and unbiased comparison across all baselines, we implemented a standardized two-stage evaluation protocol. First, we conducted automated hyper-parameter optimization using the Optuna framework [30] to identify the optimal configuration for the downstream MLP classifiers. To prevent information leakage and overfitting, this tuning phase was performed exclusively on node embeddings generated by a version of the FLASH model pre-trained on a separate, independent data split (using a random seed distinct from the three seeds used in the final evaluation). We executed 50 optimization trials for each task, searching across learning rates, dropout probabilities, number of hidden layers, hidden layer dimensions, weight decay coefficients, and patience.

Once the optimal hyperparameters were identified for a specific task (e.g., functional selectivity binary), they were fixed and applied uniformly to the MLP classifiers of all nine baselines and FLASH. For the final benchmarking, each baseline was independently pre-trained on three distinct, task-specific graph splits (defined by three random seeds). The resulting three sets of frozen embeddings per baseline were then used as inputs to the fixed-HP classifiers. This protocol ensures that any observed performance gains are attributable to the intrinsic quality and relational information captured by the pre-trained embeddings rather than task-specific fine-tuning of the downstream architecture. Task-specific MLP hyperparameters are provided in Table 18.

## 5 Supplementary Tables

**Supplementary Table 1.**
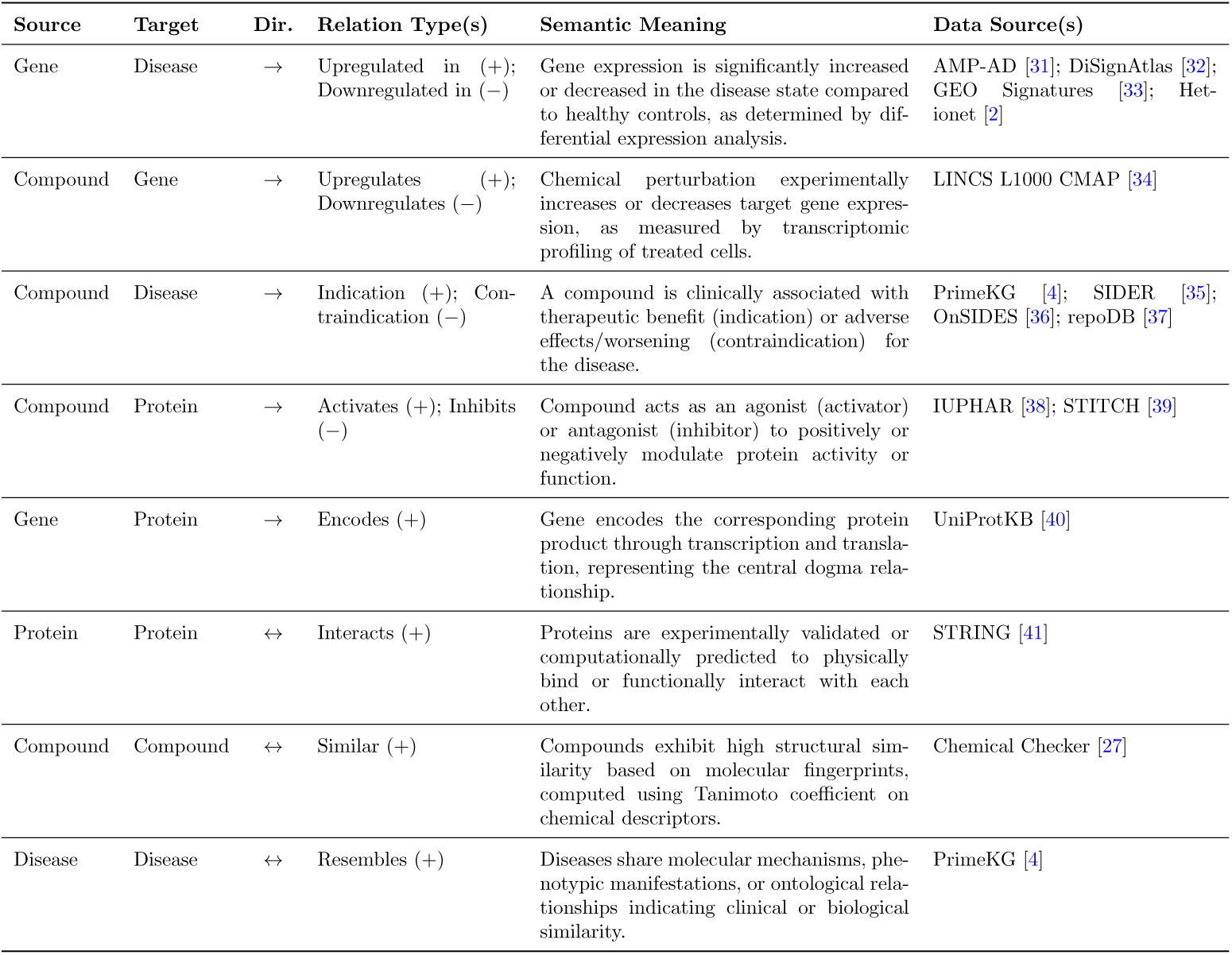
SIGMA-KG relation types and data sources. For each entity pair, we list the signed relation type(s), semantic interpretation, and contributing databases. Directed edges (→) denote asymmetric relations; bidirectional edges (↔) denote symmetric relations.

**Supplementary Table 2.** SIGMA-KG edge statistics by relation type and data source. For signed relations, positive (+) and negative (−) edge counts are reported separately. Percentages indicate the contribution of each relation type to the total knowledge graph.

| Relation Type | Data Source | Positive (+) | Negative (−) | Subtotal | Total | % |
| --- | --- | --- | --- | --- | --- | --- |
| Gene-Disease | AMP-AD [31] | 837 | 575 | 1,412 |  |  |
|  | DiSignAtlas [32] | 709,280 | 683,033 | 1,392,313 |  |  |
|  | GEO Signatures [33] | 44,491 | 44,337 | 88,828 |  |  |
|  | Hetionet [2] | 3,297 | 3,458 | 6,755 | **1,489,308** | **38.3%** |
| Compound-Gene | LINCS L1000 CMAP [34] | 669,967 | 652,863 | 1,322,830 | **1,322,830** | **34.0%** |
| compound-disease | PrimeKG (indication) [4] | 3,259 | – | 3,259 |  |  |
|  | PrimeKG (off-label) [4] | 1,556 | – | 1,556 |  |  |
|  | repoDB [37] | 7,081 | – | 7,081 |  |  |
|  | PrimeKG (contraindication) [4] | – | 20,197 | 20,197 |  |  |
|  | SIDER [35] | – | 1,908 | 1,908 |  |  |
|  | OnSIDES [36] | – | 125,956 | 125,956 | **159,957** | **4.1%** |
| compound-protein | IUPHAR [38] + STITCH [39] | 6,389 | 7,322 | 13,711 | **13,711** | **0.4%** |
| Gene-Protein | UniProtKB [40] | 19,337 | – | 19,337 | **19,337** | **0.5%** |
| Protein-Protein | STRING [41] | 251,762 | – | 251,762 | **251,762** | **6.5%** |
| Compound-Compound | Chemical Checker [27] | 581,872 | – | 581,872 | **581,872** | **15.0%** |
| Disease-Disease | PrimeKG (disease) [4] | 33,248 | – | 33,248 |  |  |
|  | PrimeKG (phenotype) [4] | 15,804 | – | 15,804 | **49,052** | **1.3%** |
| **Total** |  | **2,348,180** | **1,539,649** |  | **3,887,829** | **100%** |

**Supplementary Table 3.** Comparative analysis of signed, relational, and unsigned architectural inductive biases. This table presents a systematic inter-class comparison between the highest-performing signed models (including FLASH, SNEA, and SGCN) and the leading unsigned (e.g., DGI, GraphSAGE) and relational (e.g., HGATE, DMGI) baselines. The comparison validates the hypothesis that explicit encoding of functional polarity (±1) in the SIGMA-KG atlas provides systematic predictive gains. ”Difference % (S-U)” and ”Difference % (S-R)” quantify the performance margin of the top signed method over the top unsigned and relational alternatives, respectively. All metrics are reported as mean ± s.d. from three independent runs. Statistical significance was assessed using a two-tailed paired *t*-test; *P* -values and *t*-statistics refer to the specific inter-class comparison (Signed vs. Unsigned or Signed vs. Relational). Hierarchical significance is indicated by ^∗^*P <* 0.05 and ^∗∗^*P <* 0.01.

**Supplementary Table 4.** Comparative performance of FLASH foundation model against leading baseline models across downstream tasks. Results are presented for target-specific mode-of-action (MoA), drug-induced clinical response (DCR), and drug–drug interaction (DDI) tasks in binary, three-class, and transductive settings. For each task and metric, FLASH is benchmarked against the overall highest-performing alternative method or, in instances where FLASH establishes the state-of-the-art, the next-best performing (runner-up) model. Data are reported as mean ± s.d. derived from three independent replicates with matched random seeds. Statistical significance was determined via a two-tailed paired *t*-test; significance levels are denoted as ^∗^*P <* 0.05 and ^∗∗^*P <* 0.01. The ”Difference (%)” column indicates the relative performance gain or margin of FLASH relative to the leading baseline. Instances where the *P* -value exceeds 0.05 indicate that FLASH achieves performance comparable to the current state-of-the-art.

**Supplementary Table 5.** Binary classification performance for target-specific mode-of-action prediction (compound–protein). Each baseline model was pre-trained on SIGMA-KG with held-out compound–protein edges removed. An MLP classifier was trained to predict edge sign (activation vs. inhibition) on the held-out test set. Metrics are reported as mean ± standard deviation across three independent replicates. **Bold**: best performance; <u>underline</u>: second-best performance.

| Model | AUROC | AUPRC | Accuracy | Macro-F1 | F1<sub>Inhibition</sub> | F1<sub>Activation</sub> |
| --- | --- | --- | --- | --- | --- | --- |
| *Unsigned baselines* |  |  |  |  |  |  |
| BGRL | 0.9283 ± 0.0056 | 0.9122 ± 0.0090 | 0.8443 ± 0.0091 | 0.8441 ± 0.0089 | 0.8464 ± 0.0126 | 0.8419 ± 0.0056 |
| DGI | 0.9404 ± 0.0073 | 0.9264 ± 0.0097 | 0.8523 ± 0.0187 | 0.8520 ± 0.0185 | 0.8539 ± 0.0233 | 0.8501 ± 0.0140 |
| InfoGraph | 0.8921 ± 0.0183 | 0.8701 ± 0.0165 | 0.8064 ± 0.0179 | 0.8059 ± 0.0182 | 0.8127 ± 0.0143 | 0.7991 ± 0.0236 |
| GraphSAGE | 0.9347 ± 0.0046 | 0.9123 ± 0.0075 | 0.8627 ± 0.0112 | 0.8626 ± 0.0112 | 0.8630 ± 0.0146 | 0.8623 ± 0.0081 |
| *Signed baselines* |  |  |  |  |  |  |
| SGCN | 0.9581 ± 0.0057 | **0.9497 ± 0.0092** | 0.8931 ± 0.0099 | 0.8929 ± 0.0098 | 0.8970 ± 0.0110 | 0.8888 ± 0.0086 |
| SDGNN | 0.9391 ± 0.0031 | 0.9206 ± 0.0016 | 0.8496 ± 0.0079 | 0.8493 ± 0.0080 | 0.8421 ± 0.0085 | 0.8564 ± 0.0074 |
| SNEA | **0.9598 ± 0.0028** | <u>0.9477 ± 0.0024</u> | <u>0.8977 ± 0.0158</u> | **0.8977 ± 0.0158** | **0.8990 ± 0.0165** | **0.8964 ± 0.0151** |
| *Relational baselines* |  |  |  |  |  |  |
| DMGI | 0.9412 ± 0.0048 | 0.9252 ± 0.0014 | 0.8664 ± 0.0077 | 0.8661 ± 0.0074 | 0.8690 ± 0.0114 | 0.8632 ± 0.0052 |
| HGATE | 0.9539 ± 0.0028 | 0.9407 ± 0.0035 | 0.8885 ± 0.0069 | 0.8882 ± 0.0068 | 0.8934 ± 0.0072 | 0.8831 ± 0.0069 |
| FLASH (ours) | <u>0.9596 ± 0.0126</u> | 0.9265 ± 0.0211 | **0.8990 ± 0.0143** | <u>0.8950 ± 0.0142</u> | <u>0.8974 ± 0.0153</u> | <u>0.8925 ± 0.0131</u> |

**Supplementary Table 6.** Three-class classification performance for target-specific mode-of-action prediction (compound–protein). Each baseline model was pre-trained on SIGMA-KG with held-out compound–protein edges removed. An MLP classifier was trained to distinguish activation (+1), inhibition (−1), and non-existent edges on the held-out test set. Metrics are reported as mean ± standard deviation across three independent replicates. **Bold**: best performance; <u>underline</u>: second-best performance.

| Model | Macro AUROC | Accuracy | Macro-F1 | F1<sub>Inhibition</sub> | F1<sub>Activation</sub> | F1<sub>Non-exist.</sub> |
| --- | --- | --- | --- | --- | --- | --- |
| *Unsigned baselines* |  |  |  |  |  |  |
| BGRL | 0.9242 ± 0.0054 | 0.9157 ± 0.0048 | 0.7605 ± 0.0149 | 0.6298 ± 0.0339 | 0.6902 ± 0.0085 | 0.9615 ± 0.0024 |
| DGI | 0.9435 ± 0.0107 | 0.9110 ± 0.0060 | 0.7423 ± 0.0116 | 0.5831 ± 0.0155 | 0.6828 ± 0.0172 | 0.9609 ± 0.0044 |
| InfoGraph | 0.9291 ± 0.0083 | 0.8937 ± 0.0116 | 0.7272 ± 0.0196 | 0.5473 ± 0.0118 | 0.6849 ± 0.0419 | 0.9495 ± 0.0067 |
| GraphSAGE | 0.9282 ± 0.0112 | 0.9100 ± 0.0021 | 0.7404 ± 0.0033 | 0.6155 ± 0.0127 | 0.6486 ± 0.0065 | 0.9571 ± 0.0027 |
| *Signed baselines* |  |  |  |  |  |  |
| SGCN | 0.9421 ± 0.0065 | 0.9215 ± 0.0007 | <u>0.8070 ± 0.0018</u> | 0.7268 ± 0.0105 | **0.7323 ± 0.0139** | <u>0.9620 ± 0.0008</u> |
| SDGNN | 0.8835 ± 0.0117 | 0.9237 ± 0.0009 | 0.7897 ± 0.0031 | 0.7223 ± 0.0141 | 0.6888 ± 0.0052 | 0.9580 ± 0.0005 |
| SNEA | **0.9602 ± 0.0013** | 0.9166 ± 0.0023 | 0.7990 ± 0.0075 | 0.7311 ± 0.0228 | 0.7054 ± 0.0106 | 0.9605 ± 0.0011 |
| *Relational baselines* |  |  |  |  |  |  |
| DMGI | 0.8638 ± 0.0175 | <u>0.9245 ± 0.0029</u> | 0.7944 ± 0.0085 | <u>0.7380 ± 0.0145</u> | 0.6856 ± 0.0116 | 0.9594 ± 0.0014 |
| HGATE | 0.9154 ± 0.0043 | 0.8860 ± 0.0038 | 0.7464 ± 0.0071 | 0.6307 ± 0.0109 | 0.6710 ± 0.0093 | 0.9375 ± 0.0025 |
| FLASH (ours) | <u>0.9464 ± 0.0049</u> | **0.9289 ± 0.0010** | **0.8072 ± 0.0039** | **0.7480 ± 0.0153** | <u>0.7116 ± 0.0070</u> | **0.9621 ± 0.0003** |

**Supplementary Table 7.** Binary classification performance for drug-induced clinical response prediction (compound–disease). Each baseline model was pre-trained on SIGMA-KG with held-out compound–disease edges removed. An MLP classifier was trained to predict edge sign (indication vs. contraindication) on the held-out test set. Metrics are reported as mean ± standard deviation across three independent replicates. **Bold**: best performance; <u>underline</u>: second-best performance.

| Model | AUROC | AUPRC | Accuracy | Macro-F1 | F1<sub>Indication</sub> | F1<sub>Contraind.</sub> |
| --- | --- | --- | --- | --- | --- | --- |
| *Unsigned baselines* |  |  |  |  |  |  |
| BGRL | 0.9934 ± 0.0007 | 0.9991 ± 0.0001 | 0.9647 ± 0.0031 | 0.8931 ± 0.0072 | 0.8057 ± 0.0126 | 0.9806 ± 0.0017 |
| DGI | **0.9940 ± 0.0007** | **0.9995 ± 0.0001** | 0.9608 ± 0.0019 | 0.8840 ± 0.0049 | 0.7897 ± 0.0087 | 0.9784 ± 0.0011 |
| InfoGraph | 0.9491 ± 0.0040 | 0.9954 ± 0.0003 | 0.6893 ± 0.0374 | 0.5619 ± 0.0272 | 0.3261 ± 0.0249 | 0.7976 ± 0.0296 |
| GraphSAGE | 0.9920 ± 0.0013 | 0.9992 ± 0.0007 | 0.9562 ± 0.0031 | 0.8710 ± 0.0075 | 0.7661 ± 0.0133 | 0.9758 ± 0.0018 |
| *Signed baselines* |  |  |  |  |  |  |
| SGCN | 0.9929 ± 0.0023 | 0.9991 ± 0.0003 | 0.9863 ± 0.0002 | <u>0.9511 ± 0.0008</u> | 0.9096 ± 0.0016 | 0.9926 ± 0.0001 |
| SDGNN | 0.9923 ± 0.0016 | 0.9990 ± 0.0003 | <u>0.9874 ± 0.0005</u> | **0.9549 ± 0.0016** | **0.9166 ± 0.0030** | <u>0.9932 ± 0.0003</u> |
| SNEA | <u>0.9938 ± 0.0018</u> | <u>0.9993 ± 0.0003</u> | 0.9845 ± 0.0014 | 0.9452 ± 0.0042 | 0.8988 ± 0.0076 | 0.9916 ± 0.0008 |
| *Relational baselines* |  |  |  |  |  |  |
| DMGI | 0.9793 ± 0.0030 | 0.9976 ± 0.0003 | 0.9635 ± 0.0032 | 0.8817 ± 0.0100 | 0.7832 ± 0.0183 | 0.9801 ± 0.0018 |
| HGATE | 0.9826 ± 0.0024 | 0.9973 ± 0.0003 | 0.9742 ± 0.0008 | 0.9119 ± 0.0035 | 0.8379 ± 0.0065 | 0.9860 ± 0.0004 |
| FLASH (ours) | 0.9827 ± 0.0012 | 0.9975 ± 0.0002 | **0.9893 ± 0.0003** | 0.9386 ± 0.0010 | 0.8862 ± 0.0018 | **0.9970 ± 0.0001** |

**Supplementary Table 8.**
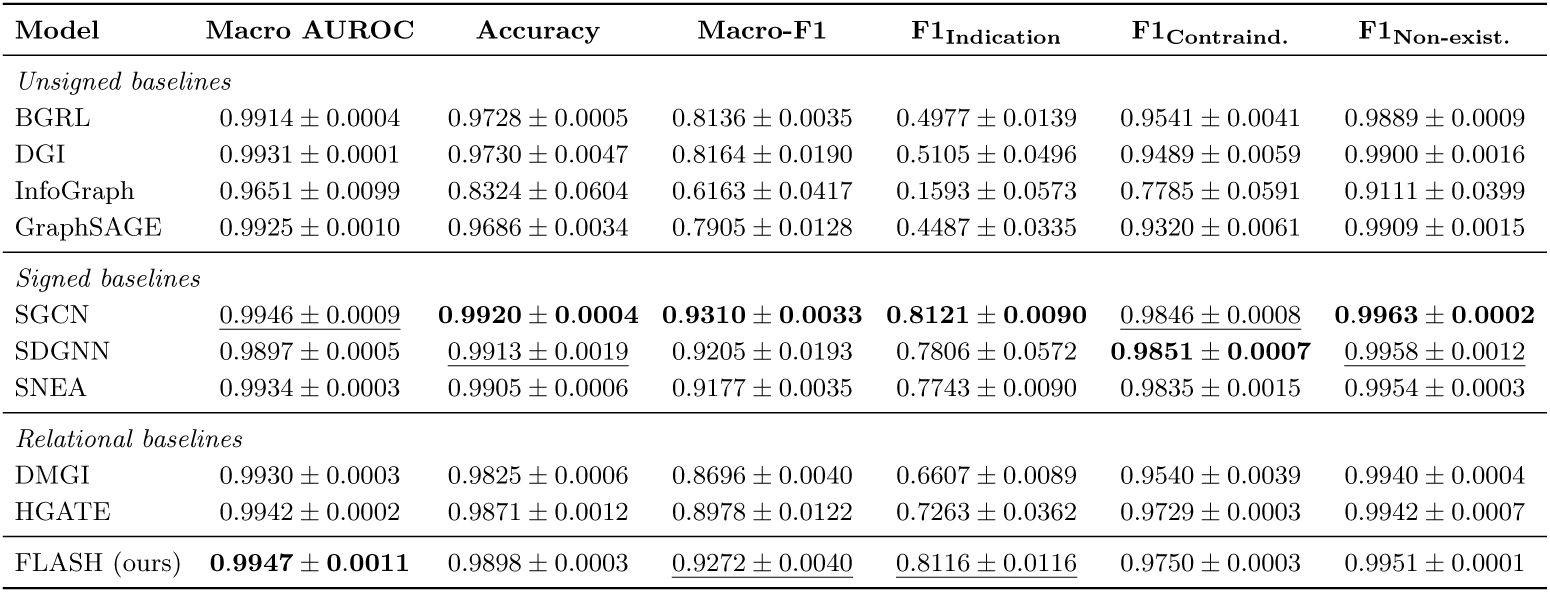
Three-class classification performance for drug-induced clinical response prediction (compound–disease). Each baseline model was pre-trained on SIGMA-KG with held-out compound–disease edges removed. An MLP classifier was trained to distinguish indication (−1), contraindication (+1), and non-existent edges on the held-out test set. Metrics are reported as mean ± standard deviation across three independent replicates. **Bold**: best performance; <u>underline</u>: second-best performance.

**Supplementary Table 9.** drug–drug interaction prediction performance on DrugBank dataset (transductive). Each baseline model was pre-trained on full SIGMA-KG (containing only structural similarity for compound-compound edges). An MLP classifier was trained to predict whether a drug pair exhibits an interaction on the external DrugBank benchmark. Metrics are reported as mean ± standard deviation across three independent replicates. **Bold**: best performance; <u>underline</u>: second-best performance.

| Model | AUROC | AUPRC | Accuracy | Macro-F1 | F1<sub>Interact.</sub> | F1<sub>Non-interact.</sub> |
| --- | --- | --- | --- | --- | --- | --- |
| *Unsigned baselines* |  |  |  |  |  |  |
| BGRL | 0.9490 ± 0.0014 | 0.9448 ± 0.0018 | 0.8774 ± 0.0024 | 0.8772 ± 0.0024 | 0.8820 ± 0.0022 | 0.8724 ± 0.0027 |
| DGI | 0.9570 ± 0.0002 | 0.9548 ± 0.0002 | 0.8873 ± 0.0013 | 0.8872 ± 0.0013 | 0.8907 ± 0.0041 | 0.8837 ± 0.0049 |
| InfoGraph | 0.8993 ± 0.0034 | 0.8871 ± 0.0055 | 0.8187 ± 0.0049 | 0.8184 ± 0.0048 | 0.8247 ± 0.0129 | 0.8129 ± 0.0131 |
| GraphSAGE | 0.9297 ± 0.0003 | 0.9232 ± 0.0004 | 0.8509 ± 0.0009 | 0.8505 ± 0.0010 | 0.8581 ± 0.0028 | 0.8429 ± 0.0036 |
| *Signed baselines* |  |  |  |  |  |  |
| SGCN | 0.9606 ± 0.0005 | 0.9584 ± 0.0006 | 0.8943 ± 0.0006 | 0.8942 ± 0.0006 | 0.8966 ± 0.0029 | 0.8918 ± 0.0031 |
| SDGNN | <u>0.9674 ± 0.0007</u> | <u>0.9666 ± 0.0008</u> | <u>0.9042 ± 0.0014</u> | <u>0.9042 ± 0.0014</u> | <u>0.9067 ± 0.0022</u> | <u>0.9017 ± 0.0026</u> |
| SNEA | 0.9568 ± 0.0008 | 0.9541 ± 0.0007 | 0.8892 ± 0.0011 | 0.8891 ± 0.0011 | 0.8925 ± 0.0015 | 0.8857 ± 0.0016 |
| *Relational baselines* |  |  |  |  |  |  |
| DMGI | 0.8443 ± 0.0299 | 0.8363 ± 0.0236 | 0.7407 ± 0.0530 | 0.7318 ± 0.0644 | 0.7182 ± 0.1132 | 0.7741 ± 0.0597 |
| HGATE | 0.9616 ± 0.0002 | 0.9598 ± 0.0002 | 0.8960 ± 0.0006 | 0.8959 ± 0.0006 | 0.8985 ± 0.0027 | 0.8933 ± 0.0031 |
| FLASH (ours) | **0.9742 ± 0.0004** | **0.9735 ± 0.0005** | **0.9166 ± 0.0014** | **0.9166 ± 0.0015** | **0.9181 ± 0.0029** | **0.9150 ± 0.0033** |

**Supplementary Table 10.**
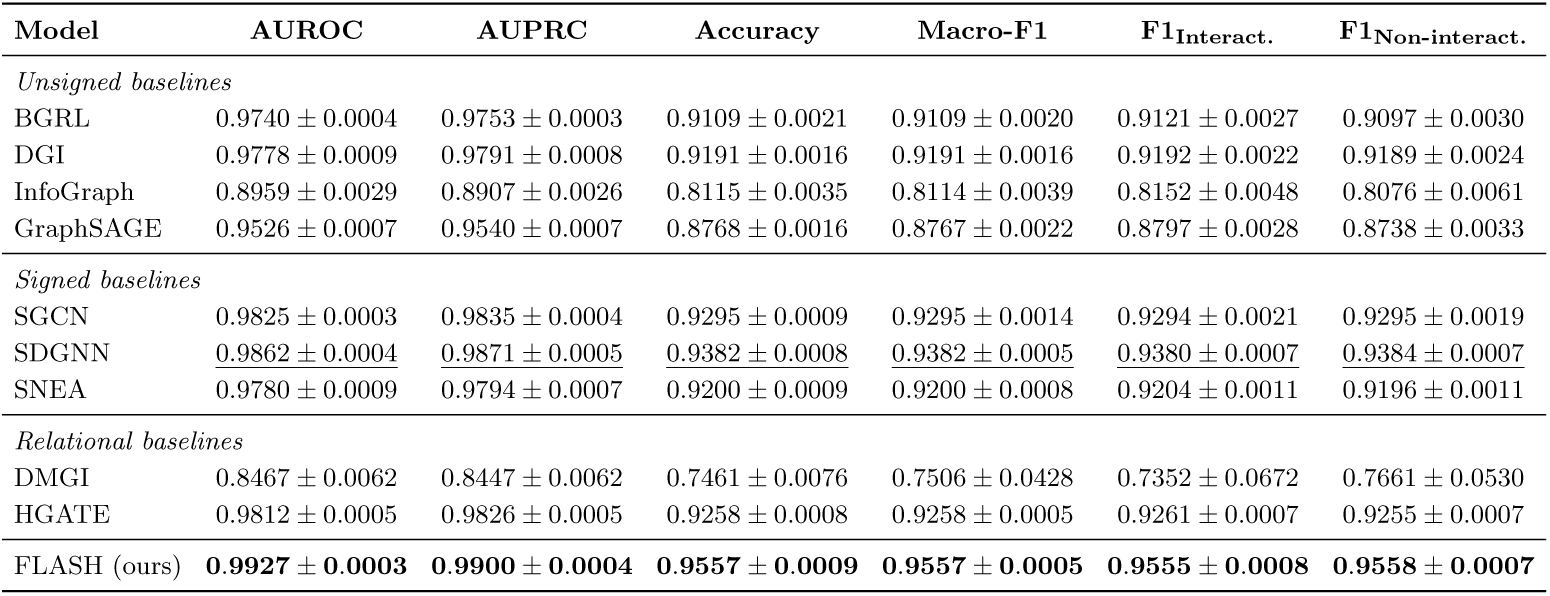
drug–drug interaction prediction performance on MUDI dataset (transductive). Each baseline model was pre-trained on full SIGMA-KG (containing only structural similarity for compound-compound edges). An MLP classifier was trained to predict whether a drug pair exhibits an interaction on the external MUDI benchmark. Metrics are reported as mean ± standard deviation across three independent replicates. **Bold**: best performance; <u>underline</u>: second-best performance.

**Supplementary Table 11.** drug–drug interaction prediction performance on DrugBank dataset (inductive).

| Model | AUROC | AUPRC | Accuracy | Macro-F1 | F1<sub>Interact.</sub> | F1<sub>Non-interact.</sub> |
| --- | --- | --- | --- | --- | --- | --- |
| *Unsigned baselines* |  |  |  |  |  |  |
| BGRL | 0.7510 ± 0.0065 | 0.7416 ± 0.0098 | 0.6799 ± 0.0074 | 0.6783 ± 0.0081 | 0.6563 ± 0.0144 | 0.7003 ± 0.0031 |
| DGI | 0.7560 ± 0.0148 | 0.7528 ± 0.0099 | 0.6883 ± 0.0082 | 0.6869 ± 0.0076 | 0.6662 ± 0.0044 | 0.7075 ± 0.0119 |
| InfoGraph | 0.7160 ± 0.0100 | 0.6977 ± 0.0133 | 0.6416 ± 0.0108 | 0.6331 ± 0.0170 | 0.5819 ± 0.0417 | 0.6842 ± 0.0099 |
| GraphSAGE | 0.7363 ± 0.0049 | 0.7265 ± 0.0025 | 0.6739 ± 0.0011 | 0.6731 ± 0.0013 | 0.6593 ± 0.0082 | 0.6870 ± 0.0063 |
| *Signed baselines* |  |  |  |  |  |  |
| SGCN | 0.7574 ± 0.0059 | 0.7539 ± 0.0138 | 0.6894 ± 0.0097 | 0.6875 ± 0.0104 | 0.6639 ± 0.0167 | 0.7110 ± 0.0074 |
| SDGNN | 0.7787 ± 0.0091 | 0.7768 ± 0.0132 | 0.7041 ± 0.0096 | 0.7022 ± 0.0100 | 0.6787 ± 0.0135 | <u>0.7258 ± 0.0072</u> |
| SNEA | 0.7781 ± 0.0102 | 0.7721 ± 0.0097 | <u>0.7069 ± 0.0107</u> | <u>0.7058 ± 0.0114</u> | <u>0.6887 ± 0.0176</u> | 0.7229 ± 0.0057 |
| *Relational baselines* |  |  |  |  |  |  |
| DMGI | 0.7406 ± 0.0185 | 0.7325 ± 0.0203 | 0.6792 ± 0.0185 | 0.6777 ± 0.0203 | 0.6671 ± 0.0392 | 0.6882 ± 0.0098 |
| HGATE | <u>0.7791 ± 0.0104</u> | <u>0.7746 ± 0.0138</u> | 0.7064 ± 0.0111 | 0.7051 ± 0.0115 | 0.6851 ± 0.0150 | 0.7250 ± 0.0082 |
| FLASH (ours) | **0.7960 ± 0.0157** | **0.7971 ± 0.0192** | **0.7189 ± 0.0154** | **0.7165 ± 0.0158** | **0.6906 ± 0.0190** | **0.7423 ± 0.0130** |

**Supplementary Table 12.** **drug–drug interaction prediction performance on MUDI dataset (inductive).**

| Model | AUROC | AUPRC | Accuracy | Macro-F1 | F1<sub>Interact.</sub> | F1<sub>Non-interact.</sub> |
| --- | --- | --- | --- | --- | --- | --- |
| *Unsigned baselines* |  |  |  |  |  |  |
| BGRL | 0.7213 ± 0.0143 | 0.7112 ± 0.0185 | 0.6615 ± 0.0148 | 0.6611 ± 0.0152 | 0.6547 ± 0.0242 | 0.6674 ± 0.0098 |
| DGI | 0.7388 ± 0.0108 | 0.7293 ± 0.0104 | 0.6725 ± 0.0083 | 0.6722 ± 0.0084 | 0.6618 ± 0.0103 | 0.6825 ± 0.0068 |
| InfoGraph | 0.7276 ± 0.0054 | 0.7193 ± 0.0039 | 0.6635 ± 0.0017 | 0.6630 ± 0.0011 | 0.6530 ± 0.0074 | 0.6729 ± 0.0094 |
| GraphSAGE | 0.7162 ± 0.0178 | 0.7075 ± 0.0184 | 0.6555 ± 0.0151 | 0.6555 ± 0.0151 | 0.6556 ± 0.0143 | 0.6554 ± 0.0164 |
| *Signed baselines* |  |  |  |  |  |  |
| SGCN | 0.7325 ± 0.0056 | 0.7190 ± 0.0050 | 0.6685 ± 0.0085 | 0.6676 ± 0.0091 | 0.6552 ± 0.0214 | 0.6800 ± 0.0043 |
| SDGNN | **0.7679 ± 0.0082** | **0.7617 ± 0.0108** | **0.6989 ± 0.0075** | <u>0.6927 ± 0.0076</u> | <u>0.6809 ± 0.0096</u> | **0.7045 ± 0.0068** |
| SNEA | 0.7493 ± 0.0069 | 0.7408 ± 0.0057 | 0.6833 ± 0.0078 | 0.6821 ± 0.0083 | 0.6638 ± 0.0141 | <u>0.7004 ± 0.0044</u> |
| *Relational baselines* |  |  |  |  |  |  |
| DMGI | 0.7163 ± 0.0199 | 0.7102 ± 0.0311 | 0.6535 ± 0.0079 | 0.6521 ± 0.0064 | 0.6665 ± 0.0216 | 0.6377 ± 0.0131 |
| HGATE | 0.7474 ± 0.0132 | 0.7283 ± 0.0153 | 0.6809 ± 0.0125 | 0.6798 ± 0.0126 | 0.6613 ± 0.0138 | 0.6983 ± 0.0121 |
| FLASH (ours) | <u>0.7595 ± 0.0064</u> | <u>0.7539 ± 0.0052</u> | <u>0.6934 ± 0.0084</u> | **0.6932 ± 0.0085** | **0.6874 ± 0.0121** | 0.6990 ± 0.0055 |

**Supplementary Table 13.**
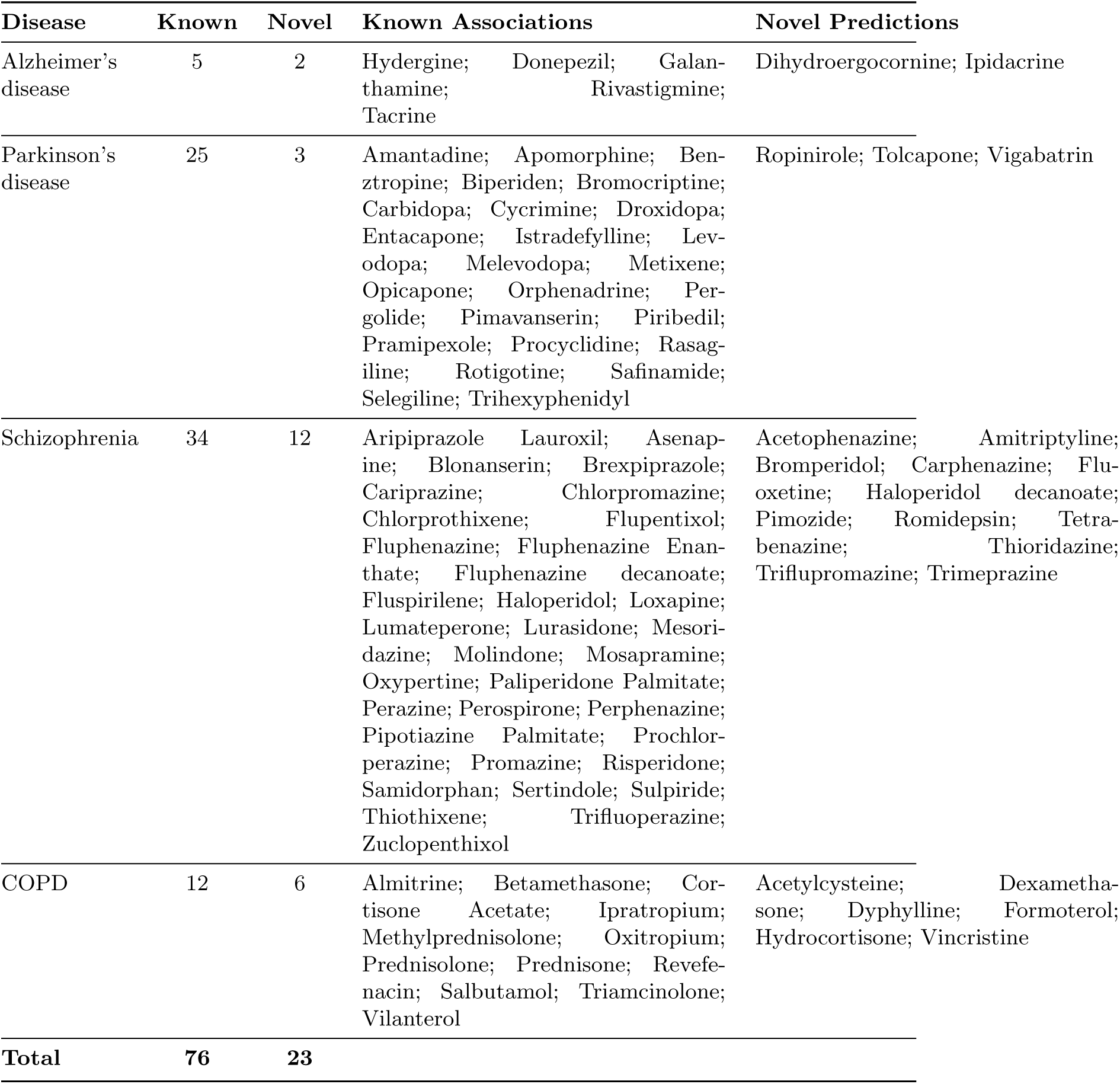
Predicted drug indications by association type. For each disease, drugs with indication probability exceeding 0.5 are categorized as: Known associations—drug-disease edges present in SIGMA-KG during pre-training, representing recovered known indications; Novel predictions—drug-disease edges absent from SIGMA-KG, representing potential repurposing candidates.

**Supplementary Table 14.**
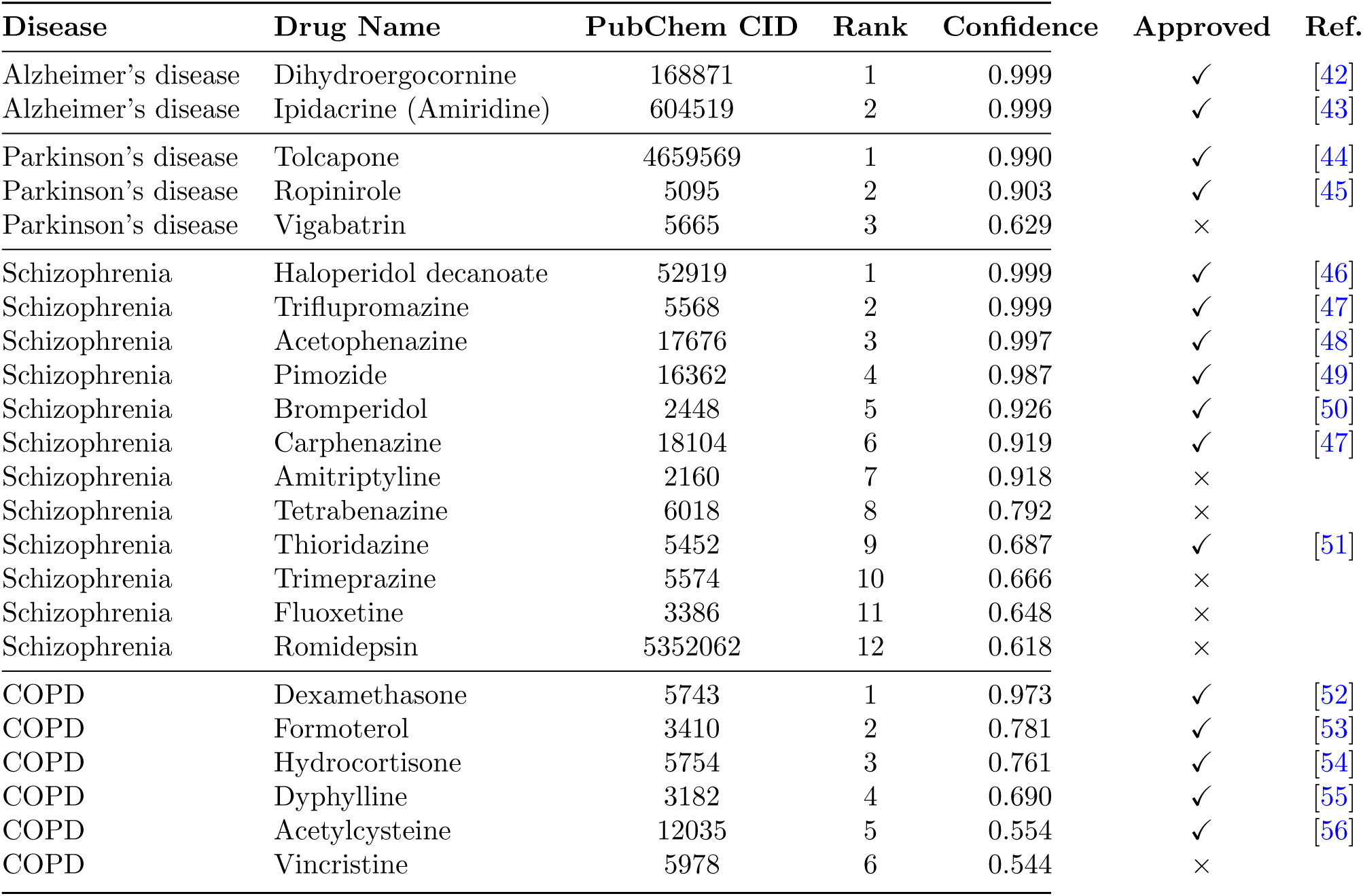
Novel drug indication predictions across five diseases. Predictions represent drug-disease associations absent from SIGMA-KG during model training. Drugs are ranked by confidence score (indication probability) within each disease. Checkmarks (✓) indicate drugs with existing regulatory approval for the predicted indication; crosses (×) indicate novel repurposing candidates without current approval for that disease.

**Supplementary Table 15.**
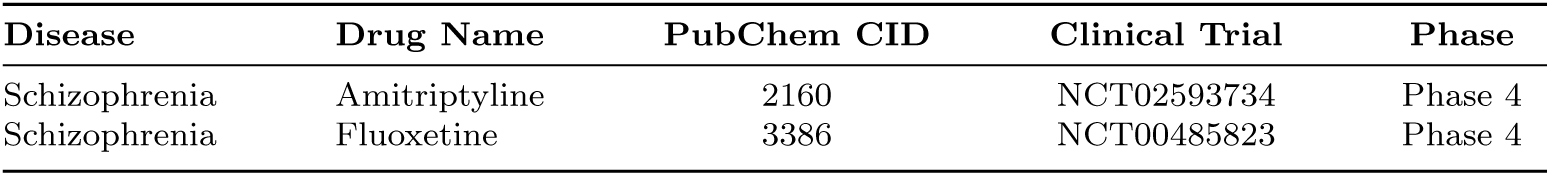
Novel indication predictions currently under clinical investigation. These drug-disease associations were predicted by FLASH as therapeutic indications and are absent from SIGMA-KG, but have active or completed clinical trials registered on ClinicalTrials.gov for the predicted disease. Note: Clinical trial identifiers correspond to registrations on ClinicalTrials.gov. Phase 4 trials are post-marketing surveillance studies conducted after regulatory approval for other indications.

| Disease | Drug Name | PubChem CID | Clinical Trial | Phase |
| --- | --- | --- | --- | --- |
| Schizophrenia | Amitriptyline | 2160 | NCT02593734 | Phase 4 |
| Schizophrenia | Fluoxetine | 3386 | NCT00485823 | Phase 4 |

**Supplementary Table 16.** Disease-specific drug targets identified through metapath analysis. For each disease, intermediate genes and proteins were aggregated across five metapaths connecting predicted drugs to their target diseases: (1) Drug–Gene–Disease, (2) Drug–Protein–Gene–Disease, (3) Drug–Drug–Gene–Disease, (4) drug–drug–Protein–Gene–Disease, and (5) drug–drug–Protein–Protein–Gene–Disease. Gene symbols follow HGNC nomenclature; protein identifiers follow UniProt accession format.

**Supplementary Table 17.** Hyperparameter configurations for self-supervised GNN baselines. All models were pre-trained using the optimal settings suggested by the original authors to ensure a fair comparison on the SIGMA-KG dataset. Key parameters represent model-specific variables critical for training stability, structural balance reasoning, or multi-relational integration.

| Model | Layers | Hidden Dim | Epochs | Learning Rate | Key Parameter (Author Suggestion) |
| --- | --- | --- | --- | --- | --- |
| BGRL | 3 | 256 | 100 | 0.01 | Feature dimension: 256 |
| DGI | 3 | 256 | 300 | 0.01 | Feature dimension: 256; Discriminator: Bilinear |
| InfoGraph | 3 | 256 | 101 | 0.01 | Feature dimension: 256; Subgraph size: 20,000 |
| GraphSAGE | 3 | 256 | 50 | 0.01 | Feature dimension: 256; Subgraph size: 20,000 |
| DMGI | 2 | 256 | 1100 | 0.001 | Feature dimension: 256; Weight decay: 0.00001 |
| HGATE | 3 | 256 | 300 | 0.0005 | Feature dimension: 256; Lambda: 0.0001; Edge sampling cap: 250,000 |
| SGCN | 2 | 256 | 222 | 0.00015 | Feature dimension: 256; Weight decay: 0.0004; Patience: 40 |
| SNEA | 2 | 256 | 271 | 0.0005 | Feature dimension: 256; Weight decay: 0.000006; Patience: 40 |
| SDGNN | 2 | 256 | 200 | 0.0001 | Feature dimension: 256; Weight decay: 0.000008; Lambda: 5; Patience: 55 |
| FLASH (ours) | 2 | 256 | 201 | 0.0002 | Feature dimension: 256; Weight decay: 0.000007; Patience: 45 |

**Supplementary Table 18.** MLP classifier architecture and hyperparameters for downstream prediction tasks. For each task, an MLP classifier was trained on pre-trained embeddings to predict edge labels. Binary tasks use ReLU (Rectified Linear Unit) activation with binary cross-entropy loss; multi-class tasks use ReLU (Rectified Linear Unit) activation with categorical cross-entropy loss. Early stopping for all classifiers (epochs=115) in this study was applied using validation-set macro-F1 and the best-performing checkpoint on the validation split was used for test evaluation.

| Task | Setting | Hidden Layers | Hidden Dims | Dropout | Learning Rate | Weight Decay | Batch Size | Patience |
| --- | --- | --- | --- | --- | --- | --- | --- | --- |
| *Functional selectivity prediction (compound–protein)* |  |  |  |  |  |  |  |  |
|  | Binary | 3 | 512 | 0.32 | 0.01 | 2e-05 | 1024 | 30 |
|  | Multi-class | 2 | 512 | 0.17 | 0.0001 | 1e-06 | 512 | 10 |
| *drug-induced clinical response prediction (compound–disease)* |  |  |  |  |  |  |  |  |
|  | Binary | 2 | 1024 | 0.15 | 0.0002 | 3e-07 | 1024 | 26 |
|  | Multi-class | 3 | 1024 | 0.30 | 0.0002 | 1e-07 | 1024 | 29 |
| *drug–drug interaction prediction* |  |  |  |  |  |  |  |  |
|  | Transductive | 3 | 1024 | 0.18 | 0.001 | 1e-05 | 512 | 21 |
|  | Inductive | 3 | 1024 | 0.32 | 0.0002 | 1e-06 | 1024 | 16 |
| *Indication prediction (Drug repurposing)* |  |  |  |  |  |  |  |  |
|  | Multi-class | 3 | 1024 | 0.25 | 0.0002 | 3e-06 | 1024 | 12 |

## 6 Supplementary Figures

**Supplementary Fig. 1.**
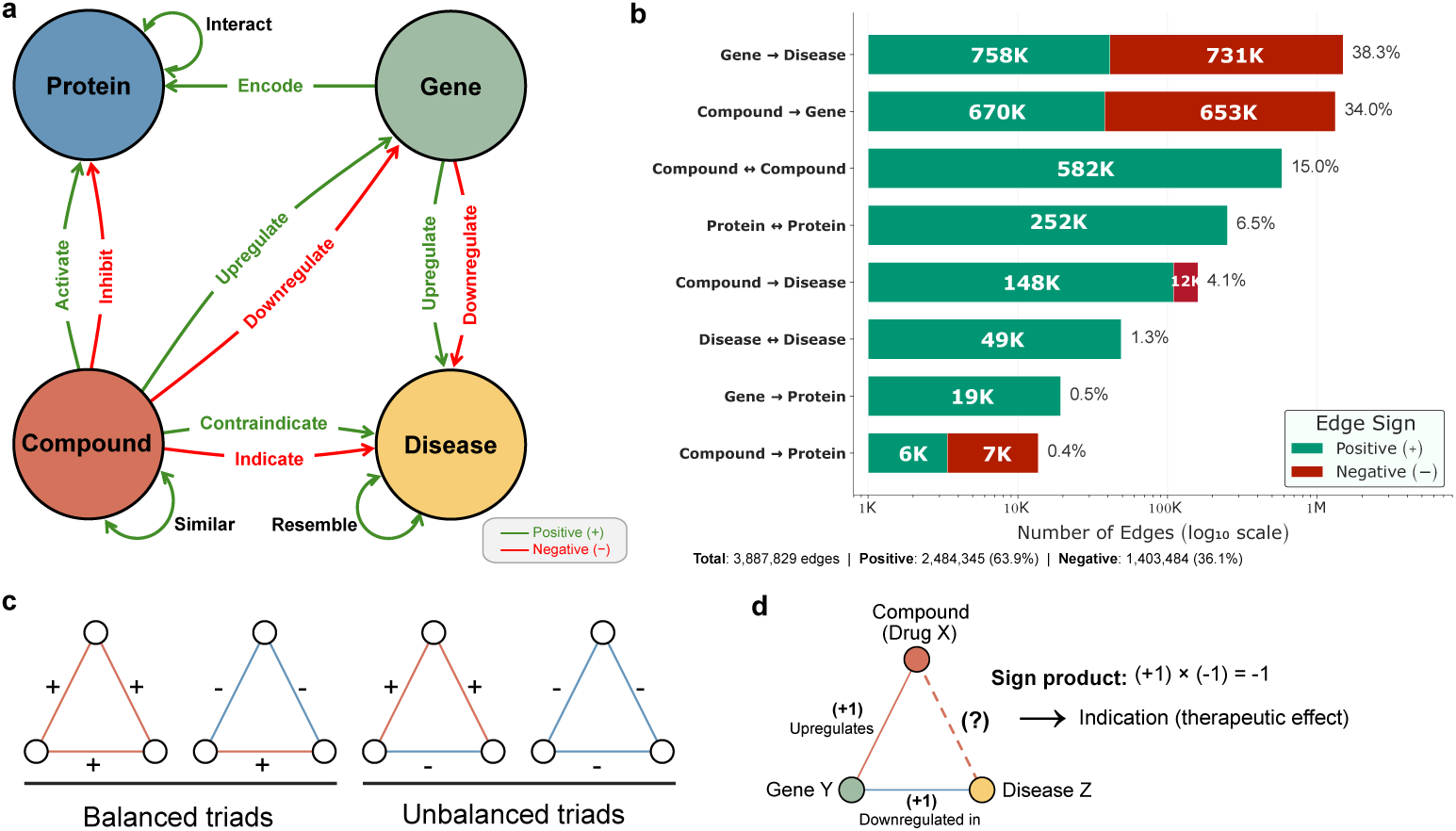
SIGMA-KG atlas schema, edge composition and sign semantics. a, Conceptual schema of the *SIGMA-KG* (SIGned Multi-omics Atlas Knowledge Graph) describing the four core entity types (compound, gene, protein and disease) and representative signed, directed relations between them. Edge signs encode mechanistic polarity (positive, negative or neutral; legend), including, for example, activation versus inhibition (compound–protein), upregulation versus downregulation (compound-gene; gene-disease), and indication versus contraindication (compound–disease), as well as unsigned similarity/relatedness links (for example, compound-compound and disease-disease). **b**, Edge-type composition of SIGMA-KG showing the number of edges per relation class (log scale) partitioned by sign (positive, blue; negative, red); percentages denote each class as a fraction of edges in the category. In total, SIGMA-KG contains 3,887,829 signed edges, comprising 2,484,345 positive (63.9%) and 1,403,484 negative (36.1%) edges. **c**, Signed triads illustrating *structural balance* (Heider’s theory): triads are balanced when the triad sign product is positive and unbalanced when the sign product is negative. **d**,Example of *transitive sign composition* along a multi-hop path: an upregulation edge (+1) from a compound to a gene composed with a downregulated-in-disease edge (−1) yields a negative sign product (−1), corresponding to a therapeutic (indication) polarity for the compound with respect to the disease in this illustrative chain (+1 × −1 = −1).

**Supplementary Fig. 2.**
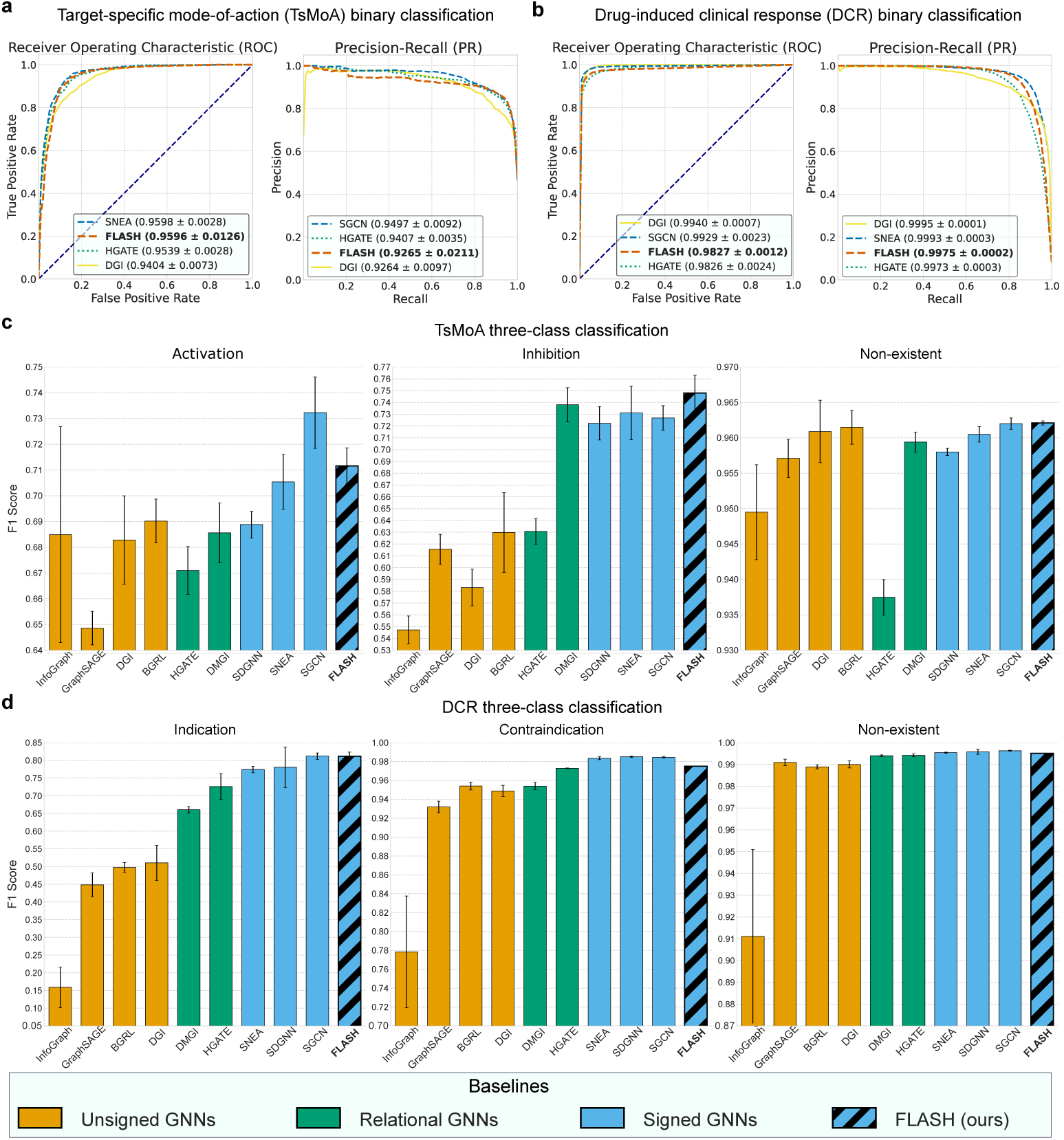
FLASH improves prediction of target-specific mode-of-action and drug-induced clinical response. a, Target-specific mode-of-action (TsMoA) *binary* classification performance shown as receiver operating characteristic (ROC) and precision-recall (PR) curves, reporting AUROC and AUPRC for FLASH and representative top baseline from each architecture paradigm. **b**, Drug-induced clinical response (DCR) *binary* classification performance shown as ROC and PR curves with AUROC and AUPRC. **c**, TsMoA *three-class* classification (activation, inhibition and non-existent) summarized by F1 score across baseline paradigms: **unsigned** GNNs (orange), **relational** GNNs (green; heterogeneous edge types without explicit sign algebra), and **signed** GNNs (blue); FLASH (ours) is shown as the hatched blue bar. **d**,DCR *three-class* classification (indication, contraindication and non-existent) summarized by F1 score using the same baseline groupings and color code. All performance values reflect results averaged over three independent runs with random train–test splits; ± values and error bars represent standard deviation.

**Supplementary Fig. 3.**
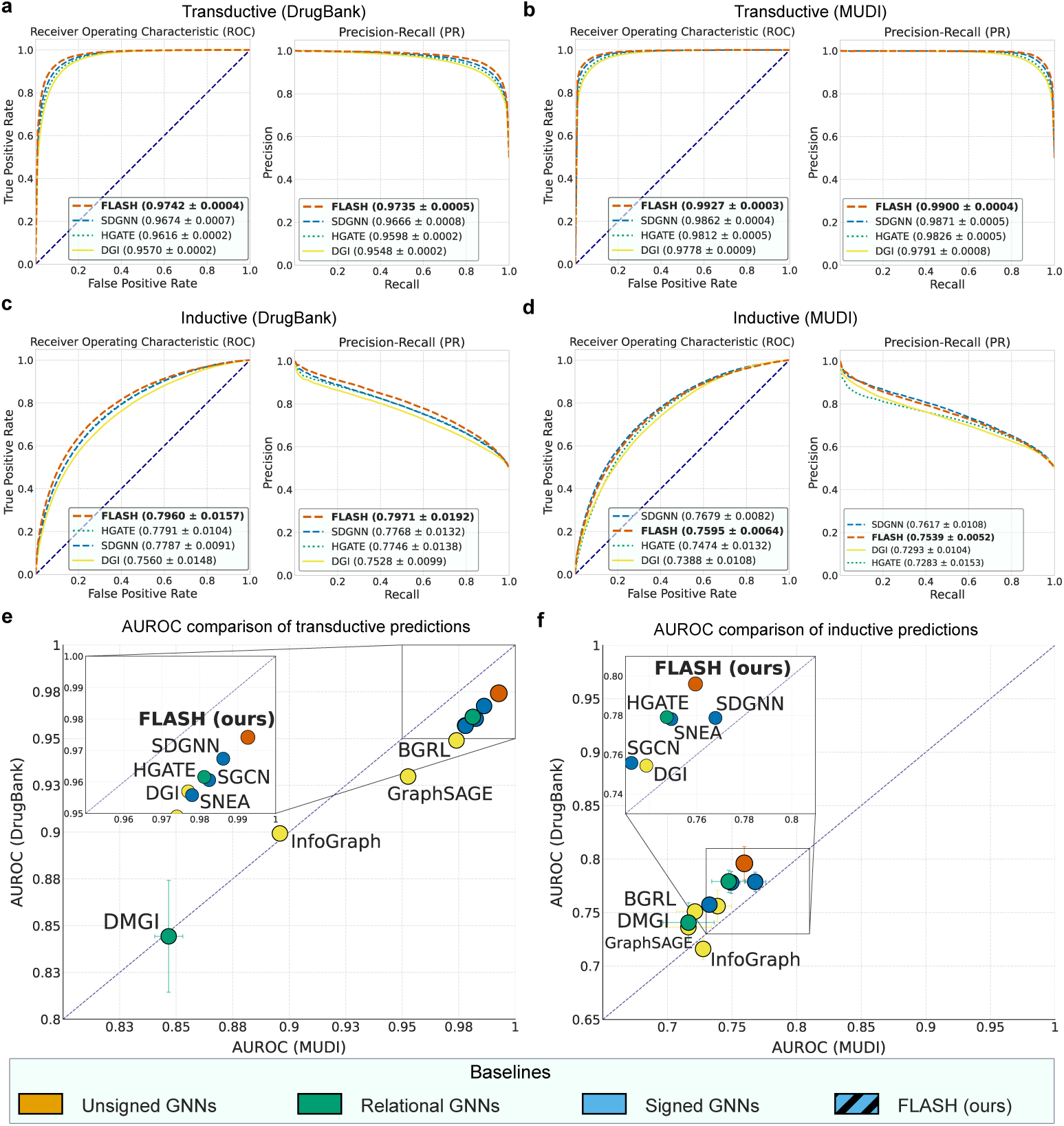
Robustness of FLASH in predicting drug–drug interactions across transductive and inductive evaluation scenarios. a,b, Receiver operating characteristic (ROC) and precision-recall (PR) curves for **transductive** DDI prediction on **DrugBank** (a) and **MUDI** (b), reporting AUROC and AUPRC for FLASH and representative top baseline from each architecture paradigm. **c,d**, ROC and PR curves for **inductive** DDI prediction on DrugBank (c) and MUDI (d), in which test drugs are unseen during training, highlighting generalization under distribution shift. **e,f**, Cross-dataset AUROC comparisons summarizing model ranking: each point corresponds to a baseline or FLASH, plotted by AUROC on MUDI (x-axis) versus DrugBank (y-axis) for transductive (e) and inductive (f) evaluations; insets magnify the high-performance region. Baseline paradigms are colour-coded as **unsigned** GNNs (orange; no sign modeling), **relational** GNNs (green; heterogeneous relations without explicit sign algebra) and **signed** GNNs (blue; explicit signed-edge modeling); **FLASH (ours)** is indicated by the hatched marker in the legend. All performance curves and values reflect results averaged over three independent runs with random train–test splits. ± values and shaded areas denote standard deviation.

**Supplementary Fig. 4.**
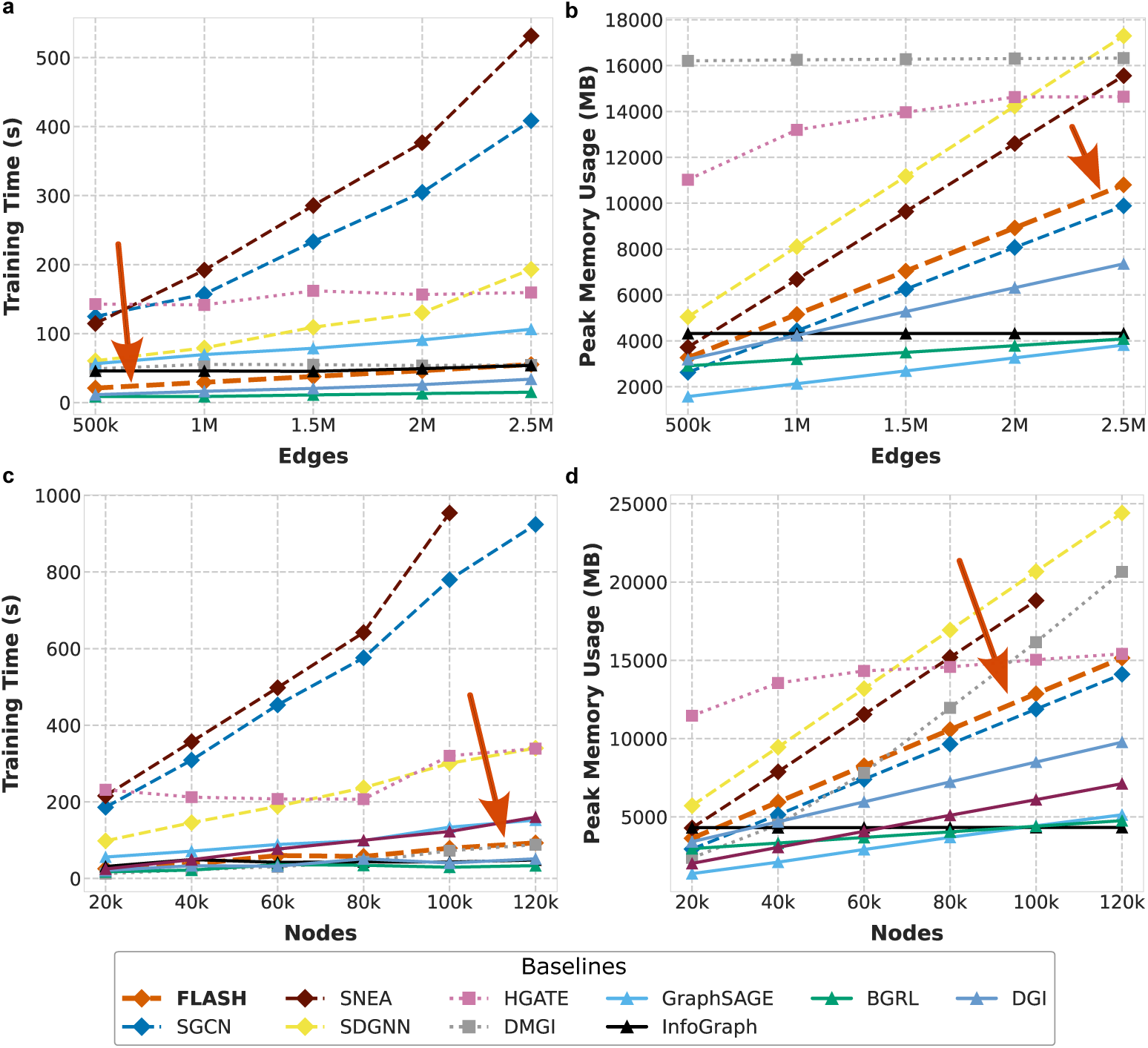
Computational efficiency and scaling of FLASH compared with baseline models across large-scale networks. a, Wall-clock training time (s) versus number of edges (500k-2.5M). **b**, Peak GPU memory (MB) versus number of edges. **c**, Wall-clock training time (s) versus number of nodes (20k-120k). **d**, Peak GPU memory (MB) versus number of nodes. *FLASH* is highlighted by an arrow in each panel. For visual disambiguation, baseline paradigms are encoded by line style/marker: **signed** baselines use dashed lines with diamond markers, **unsigned** baselines use solid lines with triangle markers, and **relational** GNNs use short-dash lines with square markers (models listed in the legend). Across all scaling regimes, FLASH achieves lower training time and/or memory usage than representative signed, relational baselines. Benchmarking was conducted on a single NVIDIA L40S GPU (48 GB). *Statistics:*none; values are single-run profiling measurements.

**Supplementary Fig. 5.**
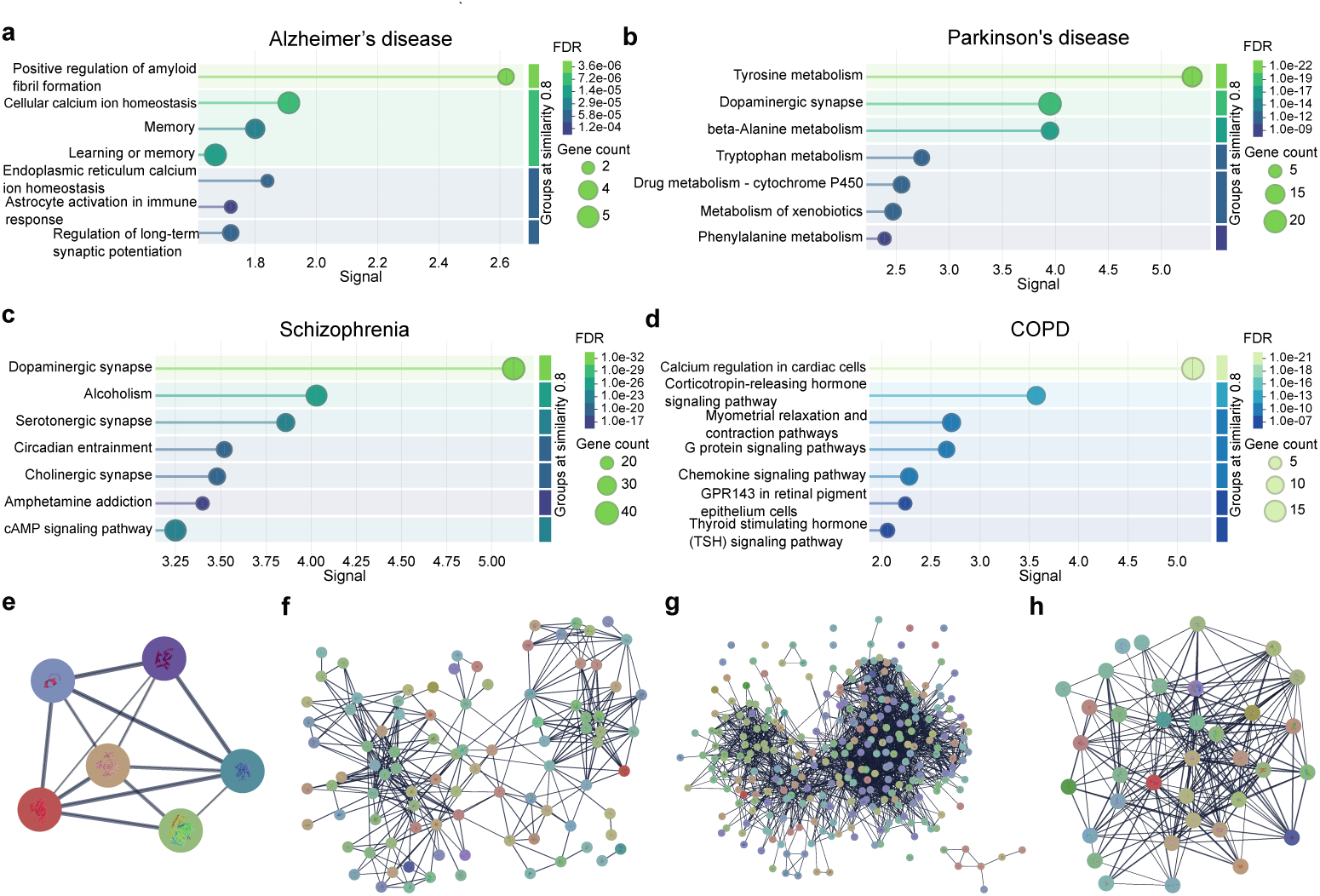
Mechanistic explainability of FLASH repurposing predictions via structurally balanced signed metapaths and disease-specific PPI modules. a-d, Pathway over-representation analysis of *mechanistic effectors* (intermediate genes and proteins) supporting FLASH-prioritized drug-disease indications (limited to novel repurposing drug candidates), obtained by tracing five predefined *multi-hop signed metapaths* in SIGMA-KG (listed in Methods) for Alzheimer’s disease (a), Parkinson’s disease (b), schizophrenia (c) and chronic obstructive pulmonary disease (COPD; d). Enrichment of disease-relevant processes (for example, amyloid fibril formation in Alzheimer’s disease and dopaminergic synapse signaling in Parkinson’s disease/schizophrenia) supports biological plausibility of the recovered cascades. Dot plots report fold enrichment (x-axis), the number of effectors mapped to each term (dot size) and Benjamini-Hochberg FDR-adjusted *P* values (colour). **e-h**, Disease-specific protein-protein interaction (PPI) modules for the same effector sets (order matched to a-d): effectors were mapped to protein identifiers and visualized as STRING interaction networks for Alzheimer’s disease (e), Parkinson’s disease (f), schizophrenia (g) and COPD (h). All modules show significantly greater connectivity than expected by chance (STRING PPI enrichment: Parkinson’s disease, schizophrenia and COPD, *P <* 1.0 × 10*^−^*^16^; Alzheimer’s disease, *P* = 0.003), consistent with coherent, system-level target modules rather than isolated proteins. *Statistics:* pathway enrichment uses a one-sided Fisher’s exact test with Benjamini-Hochberg correction (FDR*<* 0.05); STRING PPI enrichment *P* values quantify the probability of observing at least the measured number of interactions in a random network of comparable size and degree distribution.

